# Functional succinate dehydrogenase deficiency and loss of ascorbic acid transporter SLC23A1 are pathognomonic adverse features of clear cell renal cancer

**DOI:** 10.1101/2020.07.05.188433

**Authors:** Ritesh K. Aggarwal, Yiyu Zou, Rebecca A. Luchtel, Kith Pradhan, Nadia Ashai, Nandini Ramachandra, Joseph M. Albanese, Jung-in Yang, Xiaoyang Wang, Srinivas Aluri, Shanisha Gordon, Venkata Machha, Alexander Tischer, Ahmed Aboumohamed, Benjamin A. Gartrell, Sassan Hafizi, James Pullman, Niraj Shenoy

## Abstract

**Background:** Reduced succinate dehydrogenase (SDH) activity resulting in adverse succinate accumulation was previously thought to be relevant only in 0.05-0.5% of kidney cancers associated with germline *SDH* mutations (categorized ‘SDH-deficient Renal Cell Carcinoma’ in the 2016 WHO classification)

**Results:** We show that under-expression of SDH subunits resulting in accumulation of oncogenic succinate is a common feature in clear cell renal cell carcinoma (ccRCC) tumors during pathogenesis and progression, with a marked adverse impact on survival in a large cohort (n=516) of ccRCC patients. From a mechanistic standpoint, we show that von Hippel-Lindau (VHL) loss induced hypoxia-inducible factor (HIF) dependent upregulation of mir-210 in ccRCC causes direct inhibition of the *SDHD* transcript. We demonstrate that reduced expression of *SDH* subunits is associated with genome-wide increase in methylation and enhancement of epithelial mesenchymal transition (EMT) in ccRCC tumors, consistent with succinate-induced inhibition of TET activity and increase in invasiveness/ migratory ability of ccRCC cells. TET-2 inhibition-induced global regulatory DNA hypermethylation drives SDH loss-induced enrichment of EMT. *SDH* subunits under-expression had a striking association with *CDHI* (E-cadherin) loss in ccRCC tumors, in keeping with succinate-induced *CDH1* hypermethylation and under-expression in ccRCC cells. Next, in conformity with recombinant TET-2 fluorescence quenching dynamics with succinate and ascorbic acid (AA, a TET enzyme co-factor), AA treatment led to reversal of succinate-induced inhibition of TET activity, *CDH1* hypermethylation and under-expression, as well as enhanced invasiveness in ccRCC cells. Furthermore, using immunohistochemical analysis and artificial intelligence quantitation, we report that ccRCC is characterized by a marked loss of ascorbic acid transporter SLC23A1 [median percent positive cells in ccRCC primary tumors (n=104) and normal kidney cortex (n=7) was 0.7 and 32.4 respectively; p=0.0012]. Lower *SLC23A1* was associated with worse survival in ccRCC (TCGA). Lastly, intravenous AA significantly prolonged survival in a metastatic ccRCC xenograft model with increased succinate and reduced *SLC23A1* expression.

**Conclusions:** Taken together, these findings strongly indicate that functional SDH deficiency is a pathognomonic adverse feature of ccRCC (which accounts for ∼80% of all kidney cancers), and that the WHO category ‘SDH-deficient RCC’ should be re-named ‘*SDH* germline mutation-associated RCC’. Furthermore, oncogenic accumulation of succinate can be abrogated by TET modulation with AA.

**STATEMENT OF SIGNIFICANCE:** In this study, we show that under-expression of succinate dehydrogenase (SDH) subunits resulting in the accumulation of oncogenic succinate is a common, adverse, epigenetic modulating feature occurring in a vast majority of clear cell renal cell carcinoma (ccRCC), during pathogenesis and progression. Functional SDH deficiency is therefore a pathognomonic feature of ccRCC (which accounts for ∼80% of all kidney cancers), and not just limited to the 0.05-0.5% of kidney cancer patients with germline *SDH* mutations. Based on the findings reported, we propose that the ‘SDH-deficient RCC’ category in the 2016 WHO classification of kidney tumors be renamed ‘SDH germline mutation-associated RCC’. Furthermore, we demonstrate that oncogenic accumulation of succinate in ccRCC can be countered by TET modulation with ascorbic acid, and that ccRCC is characterized by a marked loss of ascorbic acid transporter SLC23A1.

Graphical abstract depicting the consequential adverse downregulation of Succinate Dehydrogenase in ccRCC and its central role in oxidative phosphorylation

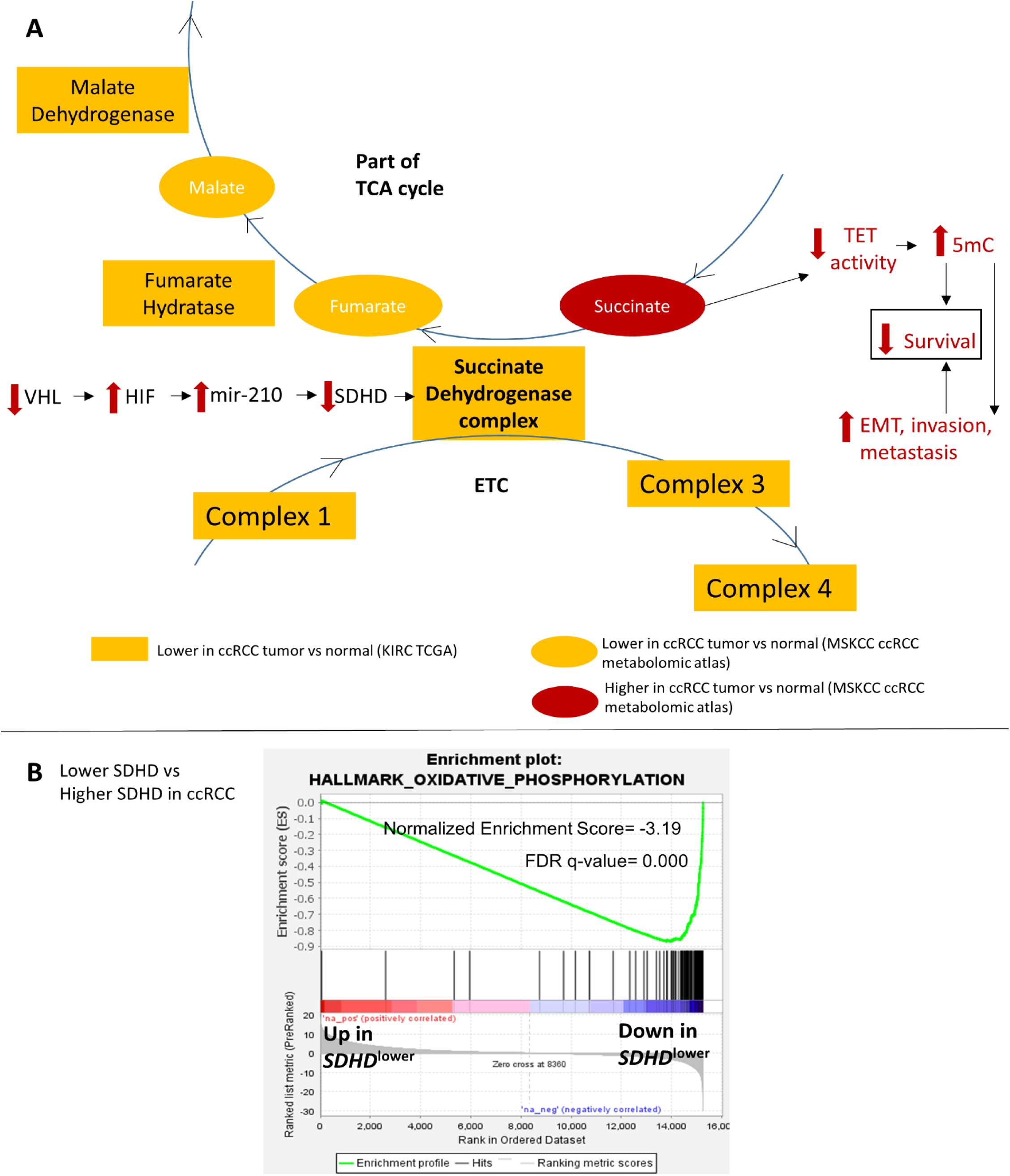

## INTRODUCTION

Clear cell renal cell carcinoma (ccRCC) is by far the most common type of kidney cancer, accounting for ∼80% of all kidney cancers ^1^. Despite recent advances, metastatic ccRCC is a generally incurable malignancy with a dismal 5-year overall survival rate of <20%, highlighting the need for further biologic and therapeutic insights in this disease.

The succinate dehydrogenase (SDH) complex is the only enzyme that is an integral component of both the TCA cycle (Krebs cycle) and the Electron Transport Chain, thus playing an important role in oxidative phosphorylation. SDH converts succinate to fumarate in the TCA cycle. The complex is located in the inner mitochondrial membrane and is composed of 4 subunits: 2 hydrophilic subunits, SDHA and SDHB; and 2 hydrophobic membrane anchoring subunits, SDHC and SDHD.

In this study, using bioinformatic analyses of ccRCC TCGA (KIRC) data (transcriptome, methylome and survival), ccRCC metabolomic repository, primary ccRCC tumors with adjacent normal renal tissue, an array of *in vitro* experiments and a metastatic ccRCC xenograft model, we-1) investigated the expression of SDH subunits in ccRCC, and the prognostic and functional consequences of loss of SDH in ccRCC; 2) shed light on the role of succinate as an important epigenetic modulating oncometabolite in ccRCC pathogenesis and progression; and 3) determined the potential of ascorbic acid (AA) in reversing the oncogenic effects of succinate in ccRCC; the expression and survival impact of AA transporters (*SLC23A1* and *SLC23A2*) in the disease as well as the efficacy of intravenous AA in improving survival in a representative metastatic ccRCC xenograft model.

## RESULTS

### Succinate dehydrogenase (SDH) subunits B, C, D are significantly under-expressed in ccRCC and associated with markedly worse survival

Analysis of the TCGA-KIRC dataset revealed that succinate dehydrogenase subunits *SDHB, SDHC* and *SDHD* are significantly down-regulated in ccRCC tumors (n=573) compared to normal renal tissue (n=72) (p<0.01 for each subunit, Figure 1A). *SDHA* expression in ccRCC, however, is not different from normal kidney tissue and does not impact survival (Figure S1).

**Fig 1:**
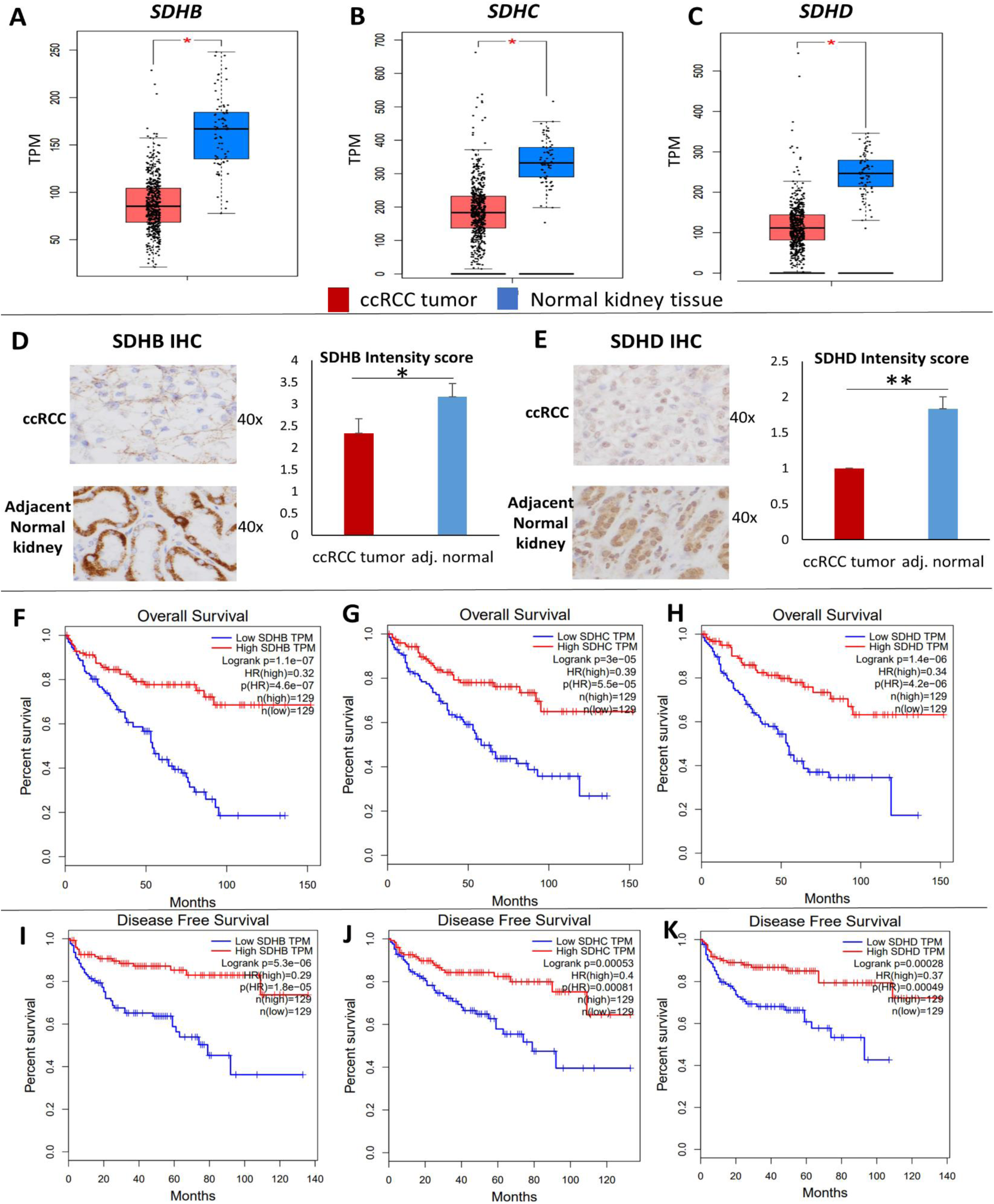
Succinate dehydrogenase subunits (B, C, D) are significantly under-expressed in ccRCC and associated with markedly worse survival. **A-C**: Analysis of the TCGA KIRC dataset revealed that succinate dehydrogenase subunits *SDHB, SDHC* and *SDHD* are significantly down-regulated in ccRCC tumor (n=573) compared to normal renal tissue (n=72) (p<0.05 for each subunit). **D, E**: SDHB and SDHD Immunohistochemistry in primary ccRCC vs adjacent normal kidney tissue revealed down-regulation of both subunits in ccRCC at the protein level. Stain intensity was on a scale of 0 (negative) to 4+ (strongly positive). For both antibodies, the non-neoplastic tubular cells in the resection stained consistently stronger than the clear cell carcinoma cells (SDHD more consistently stronger than SDHB). Stromal and other non-epithelial cells stained weakly or not at all with both antibodies (n=6 ccRCC tumors with adjacent normal, mean +/-SEM, *: p<0.05, **: p<0.01, individual cases staining depicted in Figure S2). **F-K**: Survival analyses of the TCGA KIRC dataset revealed a markedly worse Overall survival (OS) (Fig. F-H) and Disease-free survival (DFS) (Fig. I-K) with lower expression of *SDHB, SDHC* and *SDHD*. The OS Hazard Ratio (HR) for high (n=129) vs low (n=129) *SDHB, SDHC* and *SDHD* expression was 0.32, 0.39 and 0.34 respectively (high vs low quartiles, p<0.001 for each). The DFS HR for high (n=129) vs low (n=129) *SDHB, SDHC* and *SDHD* was 0.29, 0.4 and 0.37 respectively (high vs low quartiles, p<0.001 for each). These data suggest a progressive loss of *SDH* subunits as ccRCC advances.

We then investigated the protein expression levels of SDHB and SDHD in ccRCC tumors compared with adjacent normal tissue using immunohistochemistry. Consistent with the mRNA expression pattern, there was loss of both SDHB and SDHD in ccRCC compared to adjacent normal kidney tissue, as evidenced by the reduced intensity of immunostaining in ccRCC cells (n=6 ccRCC tumors with paired normed tissue, paired t-test: p<0.05 for SDHB, p<0.01 for SDHD, Figure 1B, Figure S2).

Survival analyses of the TCGA-KIRC dataset revealed a markedly worse overall survival (OS) and disease-free survival (DFS) with lower expression of *SDHB, SDHC* and *SDHD*. The OS Hazard Ratio (HR) for high (n=129) vs low (n=129) *SDHB, SDHC* and *SDHD* expression was 0.32, 0.39 and 0.34 respectively (high vs low quartiles, p<0.001 for each, Figure 1C). The DFS HR for high (n=129) vs low (n=129) *SDHB, SDHC* and *SDHD* was 0.29, 0.4 and 0.37 respectively (high vs low quartiles, p<0.001 for each, Figure 1D). Furthermore, weaker immunostaining of SDHB has been shown to be adversely prognostic in ccRCC ^2^.

### Down-regulation of SDH is a critical brake in the Krebs cycle during ccRCC pathogenesis and progression

In order to determine whether the down-regulation of SDH in ccRCC results in an accumulation of succinate, we analyzed the metabolomic repository of ccRCC ^3^. Primary ccRCC tumors (n=138) had 2-fold higher succinate compared to adjacent normal kidney tissue (n=138) (p<0.001) (Figure 2A). Furthermore, advanced (stage III/ IV) ccRCC tumors (n=90) had 1.6-fold higher succinate compared to early stage (stage I/ II) ccRCC tumors (n=48) (p=0.003) (Figure 2B).

**Fig 2:**
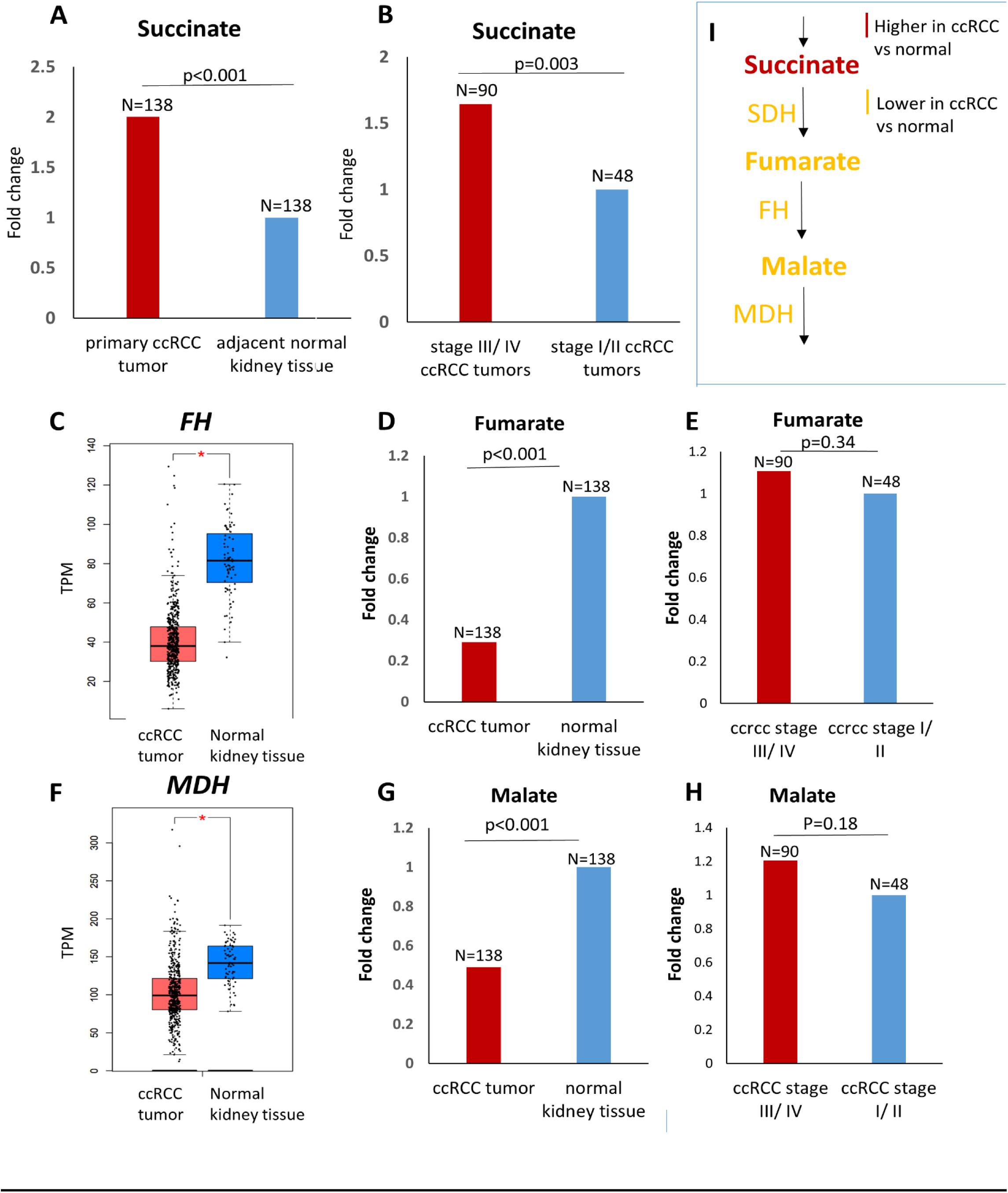
Down-regulation of SDH complex is a critical brake in Krebs cycle during ccRCC pathogenesis and progression. **A**. Analysis of the ccRCC metabolite repository revealed that primary ccRCC tumors (n=138) had 2-fold higher succinate compared to adjacent normal kidney tissue (n=138) (p<0.001). **B**. Advanced (stage III/ IV) ccRCC tumors (n=90) had 1.6-fold higher succinate compared to early stage (stage I/ II) ccRCC tumors (n=48) (p=0.003). **C, F**. Fumarate hydratase and malate dehydrogenase are significantly lower in ccRCC tumor (n=523) compared to normal renal tissue (n=72) (KIRC TCGA, p<0.01). **D, G**. Despite lower fumarate hydratase and malate dehydrogenase, fumarate and malate are significantly lower in primary ccRCC tumor (n=138) compared to adjacent normal renal tissue (n=138) (0.3-fold and 0.5-fold respectively, p<0.001 for each). **E, H**. No significant difference in the malate and fumarate content between advanced (stage III/ IV) ccRCC tumors (n=90) and early stage (stage I/ II) ccRCC tumors (n=48). **I**. Schematic representation depicting accumulation of succinate with reduced succinate dehydrogenase in ccRCC, but reduced fumarate and malate despite reduced fumarate hydratase and malate dehydrogenase, highlighting that loss of SDH is a critical brake in Krebs cycle in ccRCC. [Figures 2A, 2B, 2D, 2E, 2G, 2H have no error bars because the ccRCC metabolomic repository from which this data is derived ^3^ reports only average log2 fold change and adjusted p value for comparison between ccRCC and adjacent normal renal tissue for each metabolite]

On the other hand, fumarate was significantly lower in primary ccRCC tumor (n=138) compared to adjacent normal tissue (n=138) (0.3 fold, p<0.001, Figure 2D), despite the fact that fumarate hydratase is significantly lower in ccRCC tumor (n=523) compared to normal renal tissue (n=72) (TCGA-KIRC, p<0.01, Figure 2C). Similarly, malate was significantly lower in primary ccRCC tumor (n=138) compared to adjacent normal tissue (n=138) (0.5 fold, p<0.001, Figure 2G), despite the fact that malate dehydrogenase is significantly lower in ccRCC tumor (n=523) compared to normal renal tissue (n=72) (TCGA-KIRC, p<0.01, Figure 2F). Furthermore, there was no significant difference in the malate and fumarate content between advanced (stage III/ IV) ccRCC tumors and early stage (stage I/ II) ccRCC tumors (Figures 2E, 2H).

Put together, these data suggest that the downregulation of SDH is a critical brake in the Krebs cycle preventing the conversion of succinate to fumarate, and resulting in accumulation of succinate. This accumulation of succinate is a feature of not just ccRCC pathogenesis (higher succinate in ccRCC tumor compared to adjacent normal) but also ccRCC progression (higher succinate in advanced stage compared to early stage ccRCC).

### Von Hippel-Lindau (VHL) loss induced hypoxia-inducible factor (HIF)-dependent upregulation of miR210 in ccRCC causes direct degradation of the *SDHD* transcript

Next, we sought to investigate the mechanism of *SDH* downregulation in ccRCC. Initially, we studied the promoter CpG island (CGI) methylation of *SDH* subunit genes in ccRCC versus normal kidney (TCGA, Figure S3). Although there was a statistically significant increase in promoter CGI methylation of *SDHB, SDHD* and *SDHC*, the degree of increase in methylation was rather small, and insufficient to explain the profound downregulation of *SDH* subunits.

Therefore, we then investigated the possibility of miRNA-mediated downregulation of *SDH* subunits. mir-210, a highly conserved microRNA, is known to be induced by hypoxia, and co-ordinates various metabolic processes under hypoxic conditions. Furthermore, *SDHD* is a predicted target of this miRNA. Given the VHL-loss induced pseudo-hypoxic signature of ccRCC, we hypothesized that mir-210 may play an important pathogenic role in ccRCC and may be involved in *SDH* downregulation in this malignancy.

We found that mir-210 is the second-most upregulated miRNA in ccRCC compared to normal kidney (11-fold, p<0.001, TCGA, Figure 3A). Survival analysis revealed that higher mir210 expression is associated with significantly worse overall survival in ccRCC [high (n=131) vs low (n=133) quartiles; log-rank test p= 0.002, TCGA, Figure 3B]. In the ccRCC cell line 786-O, ChIP-seq for Hypoxia-inducible factor 2α (HIF2α) and Hypoxia-inducible factor 1β (HIF1β) revealed strong binding of both HIF2α and HIF1β to the miR210 promoter region (Figure 3C, data accessed through GSE34871 ^4^). In fact, of all the miRNAs with a binding site for HIF2α in 786-O cells, the strength of binding of HIF2α on the mir210 promoter is the highest by far (suppl. Table S1, data accessed through GSE86092 ^5^). We then mapped mir-210, *SDHD* and *SDHB* expression in ccRCC (n=507) and normal kidney (n=49) (TCGA, Figure 3D). mir-210 expression had a significant negative correlation with *SDHD* (R= -0.47, p<0.001, Figure 3E) and with *SDHB* (R= -0.38, p<0.001, Figure S4A).

**Fig 3:**
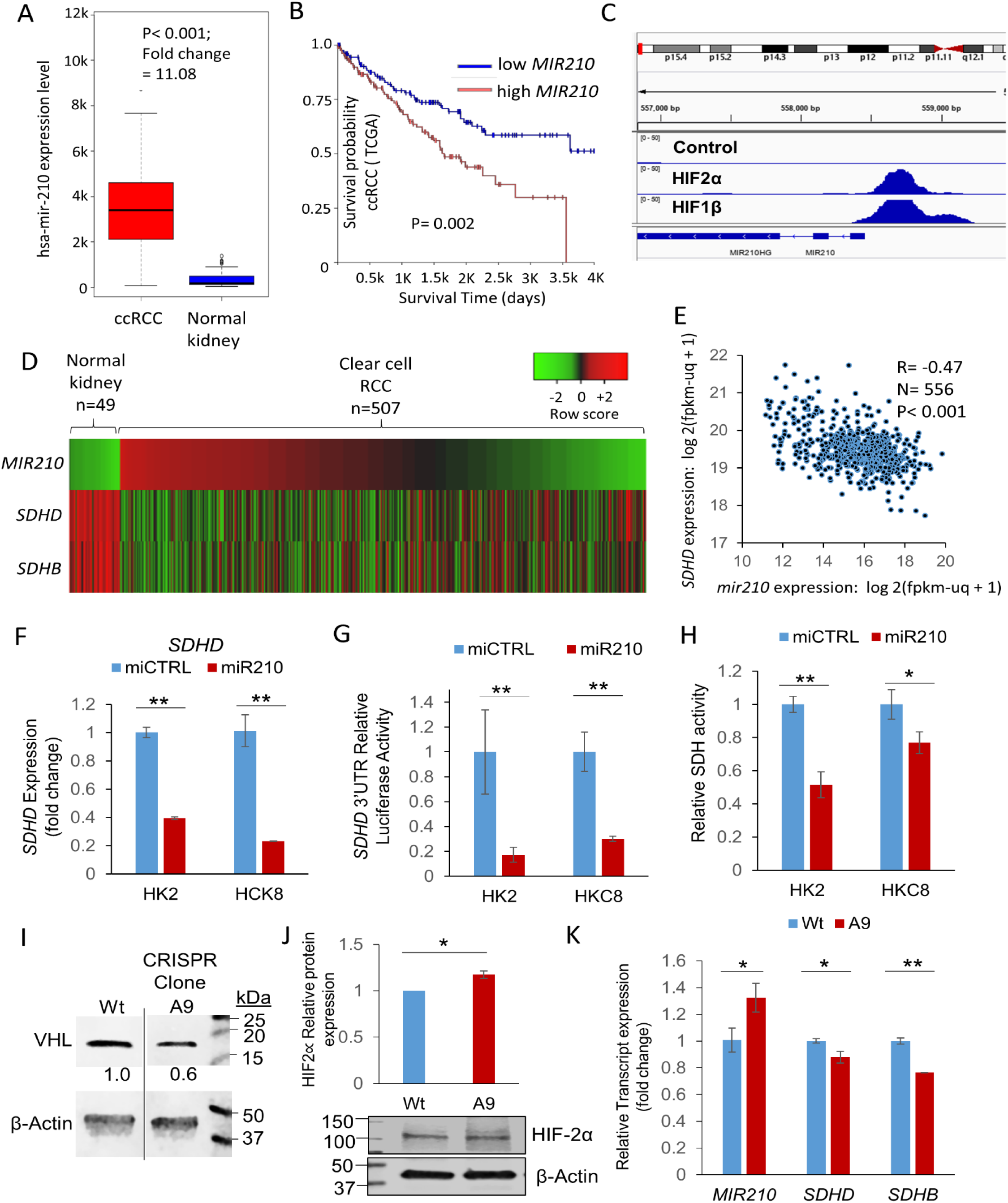
Von Hippel-Lindau (VHL) loss induced hypoxia-inducible factor (HIF) dependent upregulation of miR210 in ccRCC causes direct degradation of the *SDHD* transcript in ccRCC. **A**. mir210 is markedly up-regulated in ccRCC compared to normal kidney (11-fold, p<0.001, TCGA) **B**. Higher mir210 expression is associated with significantly worse survival in ccRCC [high vs low quartiles depicted; high (n= 131): >16.86 log 2(fpkm-uq + 1), low (n=133): <14.68 log 2(fpkm-uq + 1); log-rank test p= 0.002, TCGA] **C**. ChIP-seq for HIF2α, HIF1β, and pre-immune control in the 786O cell line. Both HIF2α and HIF1β show binding to the MIR210 promoter region. Data were accessed through GSE34871 ^4^. **D**. Marked inverse correlation between mir210 expression and SDHD expression (R=-0.47, TCGA). **E**. Heatmap showing mir210 expression along with SDHD and SDHB expression in ccRCC & normal kidney. (‘n’ as indicated, TCGA). **F**. miR210 or control miRNA were transfected into HK2 and HKC8 normal kidney cells. Cells were harvested 48 hours post-transfection for SDHD gene expression. miR210 significantly inhibited SDHD expression in both HK2 (P<0.001) and HKC8 (P=0.001) cell lines. **G**. miR210 or control miRNA were co-transfected with a SDHD 3’UTR reporter construct in HK2 and HKC8 cells, and luciferase activity was determined 48 h post-transfection. miR210 inhibited SDHD 3’UTR activity in both HK2 (P=0.004) and HKC8 (P=0.002) cell lines. **H**. miR210 or control miRNA were transfected into HK2 and HKC8 normal kidney cells. SDH activity was measured 48 hours post-transfection. miR210 inhibited SDH activity in HK2 (P=0.006) and HKC8 cell lines (P=0.05). **I**. CRISPR/Cas9 was used to generate a VHL knockdown clone, A9, from the HK2 cell line. VHL loss was validated by Western blot and Sanger sequencing (please see figure S5). **J**. Western blot analysis showed elevated HIF2α protein expression in A9 compared to Wt cells (P=0.023). **K**. Knockdown of VHL in HK2 kidney cells is sufficient to cause upregulation of hsa-miR-210-3p and downregulation of SDHD and SDHB observed in ccRCC. (F-H,J,K. All data are indicated as means ± s.e. P-values for each pairwise comparison were derived from one-sided student’s T-test. *P≤0.05; **P≤0.01)

Next, we transfected mir-210 in immortalized human kidney cell lines HK-2 and HKC-8, and found that it led to marked downregulation of *SDHD* expression in both HK2 (p<0.001) and HKC8 (p<0.001) cells (Figure 3F). However, mir-210 transfection did not decrease *SDHB* expression (Figure S4B).

In order to determine whether *SDHD* downregulation by mir-210 in these cell lines was through direct degradation of the *SDHD* transcript or an indirect mechanism, we co-transfected mir-210 (or control miRNA) with an SDHD 3’UTR reporter construct in HK2 and HKC8 cells, and luciferase activity was determined 48 h post-transfection. mir-210 inhibited *SDHD* 3’UTR activity in both HK2 (P=0.004) and HKC8 (P=0.002) cell lines (Figure 3G), establishing that mir-210 directly degrades *SDHD*.

Next, we aimed to determine whether mir-210 decreased total SDH enzymatic activity. SDH activity was measured 48 hours post-transfection of mir-210 or control miRNA into HK2 and HKC8 cells. As hypothesized, mir-210 inhibited SDH activity in both HK2 (P=0.006) and HKC8 (P=0.05) cell lines (Figure 3H).

Finally, we aimed to determine whether loss of VHL was sufficient to induce upregulation of mir-210 and downregulation of *SDH* subunits. CRISPR mediated monoallelic deletion of *VHL* (Figure 3I, Figure S5) in the HK2 cell line caused an increase in HIF2α expression (Figure 3J), which was concomitant with increased mir-210 and decreased *SDHD/ SDHB* expression (Figure 3K). (*VHL* deletion confirmed with Sanger Sequencing-Figure S5).

In keeping with the findings above, in the ccRCC cell line 786-O cultured under long-term hypoxic conditions (1% O_2_, 3 months), both an increased mir-210 expression as well as decreased *SDHD*/ *SDHB* expression were observed (Figure S6, data accessed through GSE107848 ^6^).

### SDH under-expression is associated with enrichment of epithelial mesenchymal transition (EMT) in ccRCC tumors; succinate increases invasiveness and migration ability of ccRCC cells

Having determined that under-expression of *SDH* subunits is associated with markedly worse prognosis, we sought to determine the key oncogenic pathway/s enriched with the loss of *SDH*. We analyzed the TCGA-KIRC dataset with the Gene Set Enrichment Analysis (GSEA) software, and found that lower *SDHD* is associated with marked enrichment of the epithelial mesenchymal transition (EMT) pathway, which aids invasion and metastasis (Normalized Enrichment Score: 2.17, Nominal p-value: 0.0, FDR q value: 0.0; Figure 4A). In fact, EMT was the most positively enriched pathway in ccRCC tumors associated with lower *SDHD* expression.

**Fig 4:**
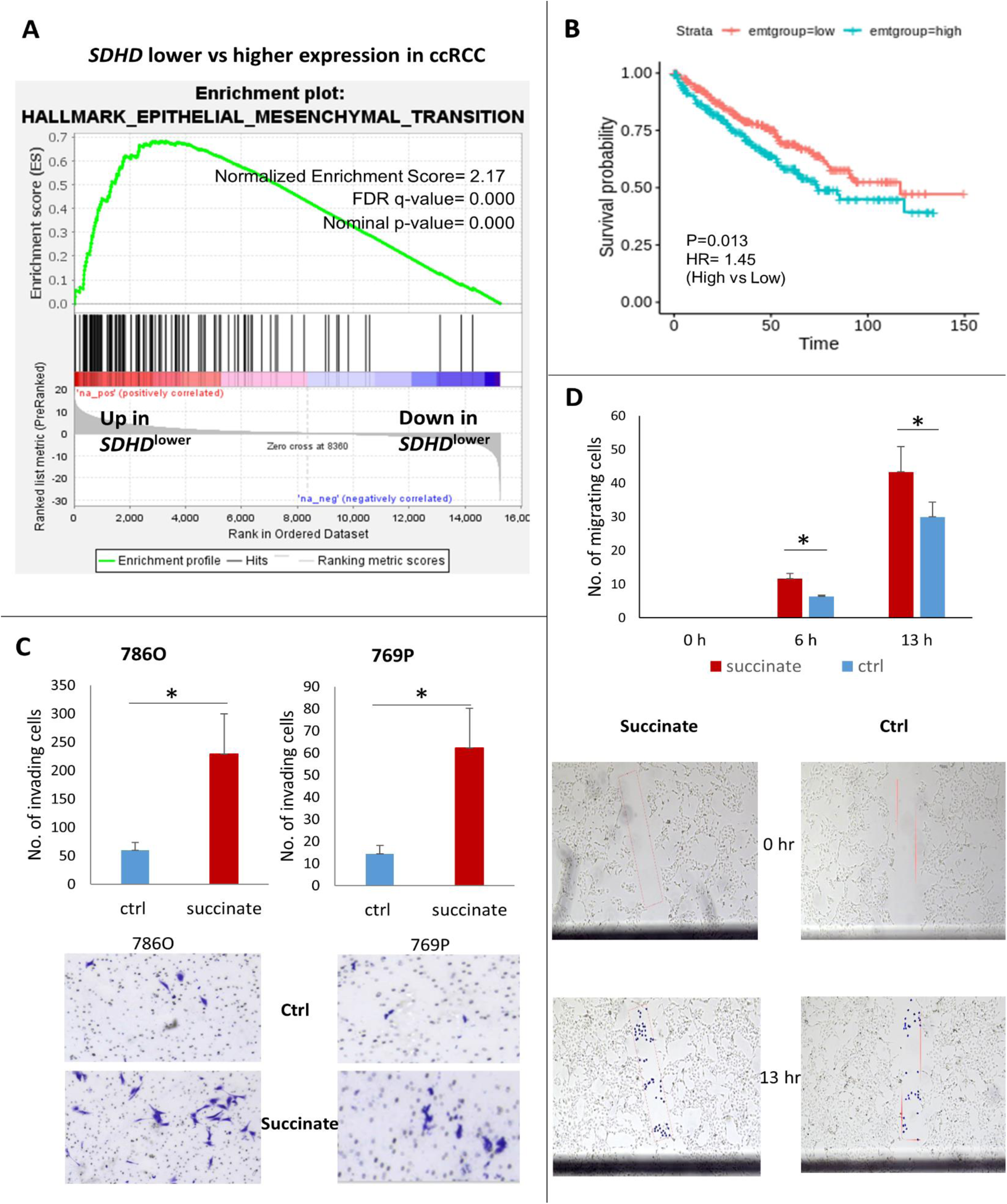
SDH under-expression is associated with enrichment of epithelial mesenchymal transition in ccRCC tumors; succinate increases invasiveness and migration ability in ccRCC cells. **A**. Analysis of the KIRC TCGA with the Gene Set Enrichment Analysis (GSEA) revealed that lower *SDHD* is associated with marked enrichment of the Epithelial Mesenchymal Transition Pathway, known to aid invasion and metastasis. **B**. Higher EMT Score is associated with worse prognosis in ccRCC (high vs low EMT Score HR: 1.46, p=0.013, KIRC TCGA). EMT Score was derived from assigning a positive score to normalized expression values of known mesenchymal markers (*FN1, VIM, ZEB1, ZEB2, TWIST1, TWIST2, SNAI1, SNA2, CDH2*) and a negative score to normalized expression values of known epithelial markers (*CLDN4, CLDN7, TJP3, MUC1, CDH1*). **C**. Succinate treatment (50uM) of ccRCC cell-lines 786-O and 769-P resulted in a significant increase in the invasiveness of ccRCC cells as determined by the Matrigel invasion assay (72-hour time-point, n=3 for each cell line, data representing mean +/-SEM, *: p<0.05). Representative pictures shown. BD BioCoat Matrigel Invasion Chambers were used to assess the invasiveness of tumor cells, with control or succinate (50uM). A thin layer of Matrigel matrix at the bottom of each chamber serves as a reconstituted basement membrane in vitro, while the chemoattractant is present in the culture medium on the outside of the chamber. Manufacturer’s protocol to prepare the matrigel chambers was followed and same number of cells per chamber (at least 20,000 cells) were plated in replicates. Incubation time of the cells in the matrigel chambers was 72 h after which the bottom of the chambers was fixed with buffered formaldehyde, stained with bromophenol blue. The number of invading cells were counted after removal of matrigel layer and reported as mean +/-SEM. **D**. The Scratch test revealed a significant increase in the migration of ccRCC cells with succinate (50uM) treatment (n=2, data representing mean +/-SEM, *: p<0.05). Representative pictures shown for 0, 6 and 13 hr timepoints. A rectangular area within the scratch (pink mark) was pre-defined for both succinate and control groups. During analysis, each cell that had invaded into the rectangular area was highlighted with a blue dot using the ImageJ imaging software, and counted digitally.

Using a previously defined EMT score ^7^, in which normalized expression values of mesenchymal markers (*FN1, VIM, ZEB1, ZEB2, TWIST1, TWIST2, SNAI1, SNA2, CDH2*) are given a positive score and that of epithelial markers (*CLDN4, CLDN7, TJP3, MUC1, CDH1*) are given a negative score, we showed that a higher EMT score is associated with worse prognosis in ccRCC with a hazard ratio of 1.46 between higher EMT score and lower EMT score (Figure 4B).

Having observed the marked enrichment of EMT pathway with lower *SDH*, we hypothesized that succinate accumulation from loss of SDH induces EMT phenotypic changes in RCC cells, aiding invasion, migration and metastasis. Indeed, exogenous succinate treatment (50uM) of ccRCC cell lines 786-O and 769-P resulted in a significant increase in the invasiveness as determined by the Matrigel invasion assay (72-hour time-point, Figure 4C). Furthermore, a Scratch assay also revealed a significant increase in the migration of ccRCC cells with succinate (50uM) treatment (72-hour time-point, Figure 4D).

### *SDHB/ SDHD* under-expression is associated with genome wide increase in cytosine methylation in ccRCC tumors driven by succinate induced inhibition of TET activity

We then aimed to determine the association between loss of SDH subunits and global cytosine methylation in ccRCC, as we have previously shown that higher methylation and lower hydroxymethylation are associated with worse prognosis in ccRCC ^8,9^. Analysis of 215 TCGA-KIRC cases with available methylation beta values and gene expression, revealed that lower *SDHD* expression (n=108) was associated with higher global cytosine methylation compared to higher *SDHD* expression (n=107) (p=0.007, Figure 5A). Similarly, lower *SDHB* expression was associated with higher global cytosine methylation compared to higher *SDHB* expression (n=107) (p=0.004, Figure 5B).

**Fig 5:**
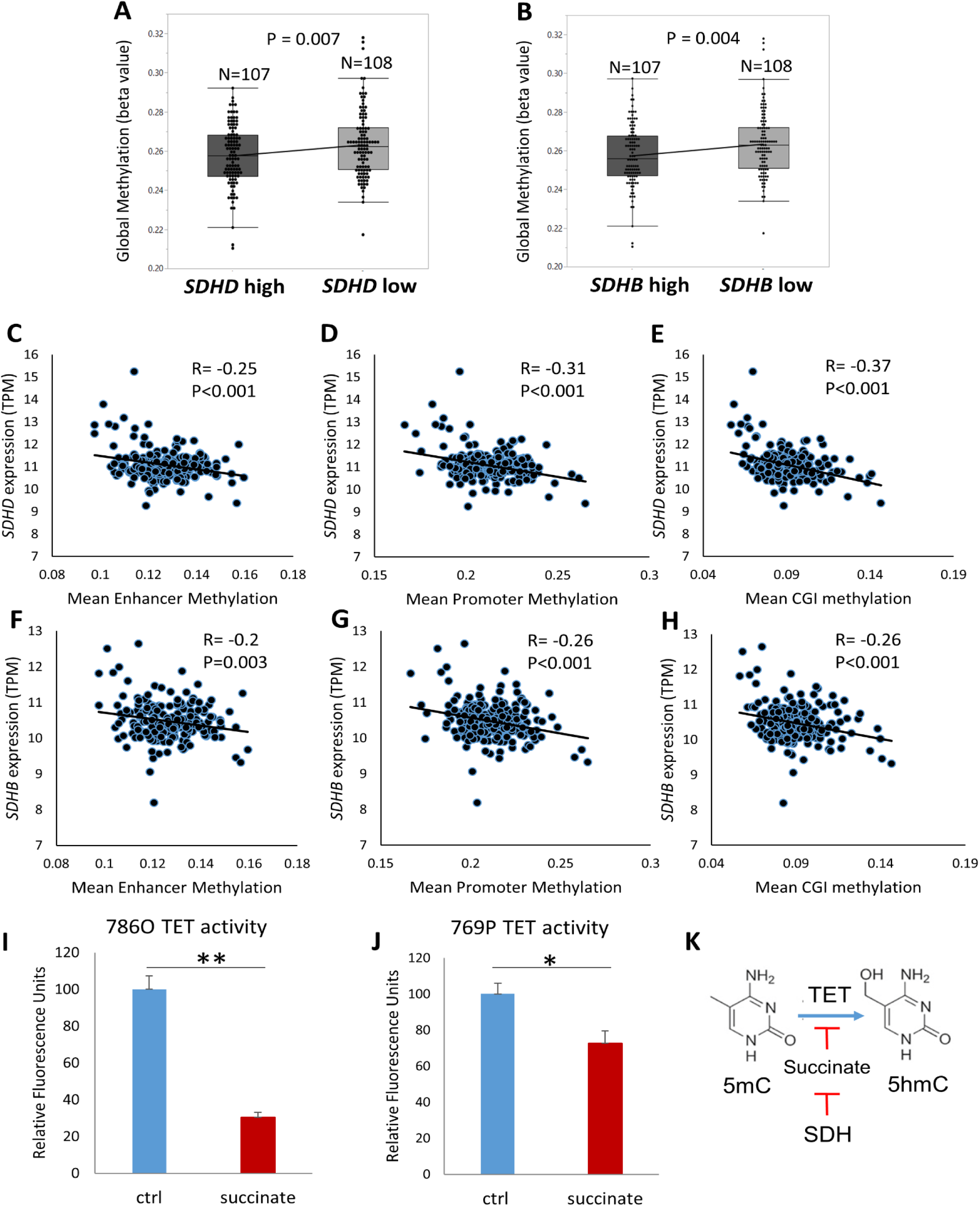
*SDHD*/ *SDHB* under-expression is associated with genome wide increase in cytosine methylation in ccRCC tumors likely driven by succinate-induced inhibition of TET activity. **A**. Analysis of 215 KIRC TCGA cases with available methylation beta values and gene expression revealed that lower *SDHD* expression (n=108) was associated with higher global cytosine methylation (mean beta values) compared to higher *SDHB* expression (n=107) (p=0.007). **B**. Similarly, lower *SDHB* expression was associated with higher global cytosine methylation compared to higher *SDHB* expression (n=107) (p=0.004). **C-E**. Analysis of specific regulatory regions of the genome revealed that *SDHD* expression had a significant inverse correlation with Enhancer methylation (R= -0.25, p<0.001), Promoter methylation (R= -0.31, p<0.001), and CpG island methylation (R= -0.37, p<0.001). **F-H**. Similarly, *SDHB* expression too had a significant inverse correlation with Enhancer methylation (R= -0.2, p=0.003), Promoter methylation (R= -0.26, p<0.001), and CpG island methylation (R= -0.26, p<0.001). **I, J**. Succinate treatment (50uM) of ccRCC cell-lines 786-O and 769-P for 24 hours resulted in significant inhibition of TET enzyme activity (n=2, *: p<0.05, **: p<0.01, ELISA based Epigentek TET activity). **K**. Schematic representation showing regulation (inhibition) of TET activity by succinate, which in turn is regulated by succinate dehydrogenase.

Having determined the inverse relation between *SDHB/ SDHD* expression with global cytosine methylation, we sought to correlate *SDHB/ SDHD* expression with individual key regulatory regions of the genome-CpG islands (CGIs), promoters and enhancers. *SDHD* expression had a significant inverse correlation with enhancer methylation (R= -0.25, p<0.001), promoter methylation (R= -0.31, p<0.001), and CGI methylation (R= -0.37, p<0.001) (Figures 5C-5E). Similarly, *SDHB* expression too had a significant inverse correlation with enhancer methylation (R= -0.2, p=0.003), promoter methylation (R= -0.26, p<0.001), and CGI methylation (R= -0.26, p<0.001) (Figures 5F-5H).

We hypothesized that the inverse correlation between SDH subunits expression and global cytosine methylation is due to the accumulation of succinate secondary to loss of SDH in ccRCC (seen in Fig 2) and the inhibitory effect of succinate on Ten-Eleven Translocation (TET) function. Succinate has been shown to be an inhibitor of multiple α-kg dependent dioxygenases, including the TET enzymes ^10^. Using an ELISA-based TET activity assay, we demonstrated that succinate treatment (50uM) of ccRCC cell lines 786-O and 769-P resulted in significant inhibition of TET enzymatic activity (Figures 5I-5K).

Similar to *SDHB/ SDHD* expression, *TET2* expression was also found to inversely correlate with global methylation levels in ccRCC. Lower *TET2* expression (n=108) was associated with higher global cytosine methylation (mean beta value) compared to higher *TET2* expression (n=107) (p=0.035, Figure S7A). Furthermore, lower *TET2* expression was associated with worse overall survival in ccRCC. The OS HR for high (n=129) vs low (n=129) *TET2* expression was 0.4 (TCGA-KIRC, high vs low quartiles, p<0.001, Figure S7B). These data strongly support a tumor suppressor role for *TET2* in ccRCC, and further highlight the adverse implications of TET-2 inhibition in this disease. In fact, only *TET2*, not *TET1* or *TET3*, is negatively correlated with global methylation in ccRCC as well as significantly associated with adverse outcome (TCGA; Figure S8).

Furthermore, stable knockdown of *SDHB* in human embryonic kidney cells (HEK293T) has been shown to markedly reduce ectopic TET-2 enzyme catalyzed 5hmC production by nearly 70% ^10^.

### TET-2 inhibition-induced global regulatory DNA hypermethylation drives SDH loss-induced enrichment of EMT, with reversal by a TET enzyme-activating demethylating agent, ascorbic acid (AA)

Next, we sought to investigate the mechanism by which loss of SDH enhances EMT. We hypothesized that global methylation changes induced by succinate accumulation results in the enrichment of EMT pathway, primarily by suppression of ‘epithelial marks’. Indeed, promoter CGI methylation and enhancer methylation in ccRCC strongly correlated with enrichment of the EMT pathway (NES-2.11 and 1.5 respectively), providing a link between SDH loss-induced genome-wide methylation and enhancement of EMT in ccRCC tumors (n=215, Figures 6A, 6B).

**Fig 6:**
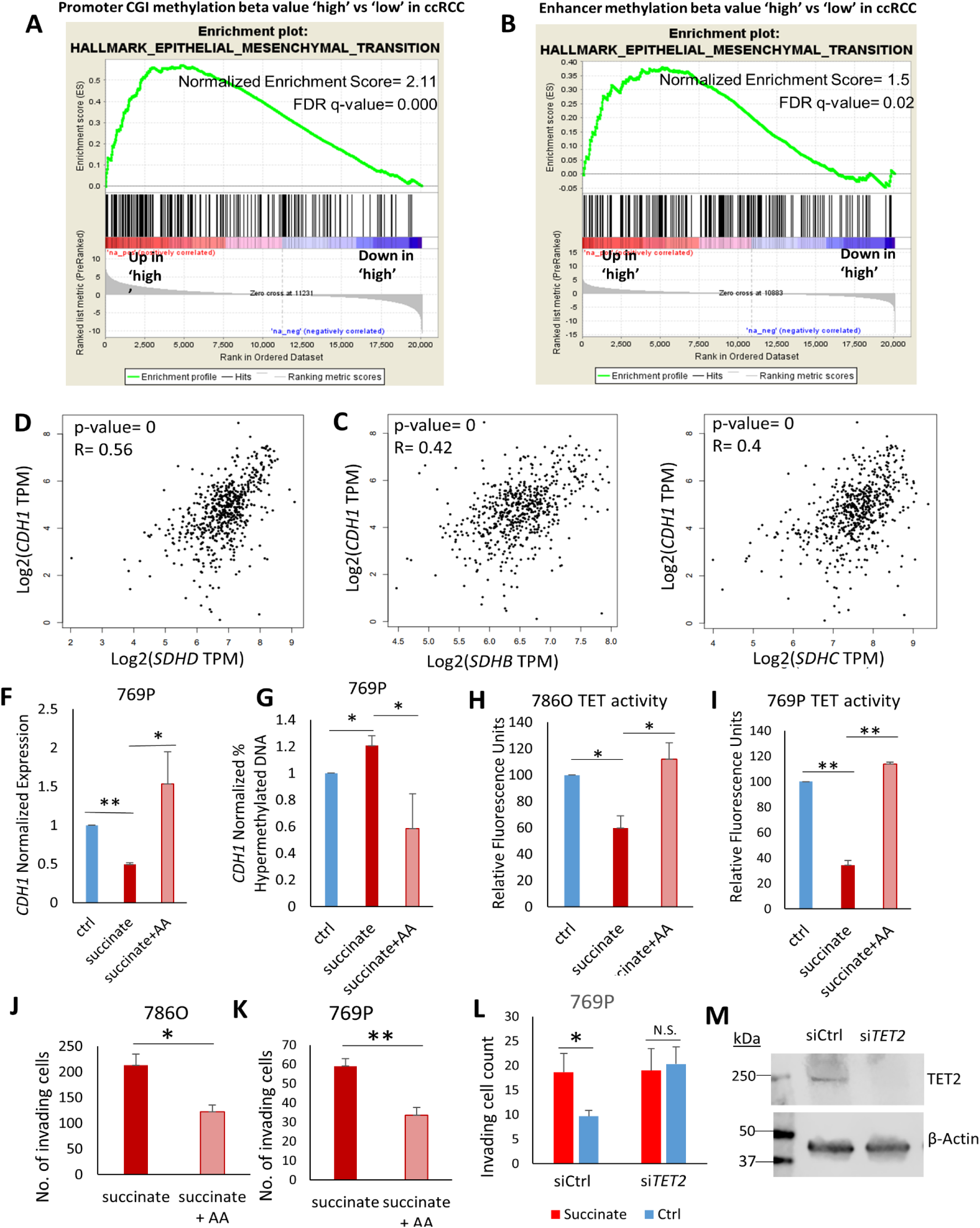
TET-2 inhibition-induced global regulatory DNA hypermethylation drives SDH loss-induced enrichment of EMT, with reversal by a TET enzyme-activating demethylating agent, ascorbic acid (AA) **A-B** Pathway enrichment analysis revealed that Promoter CGI methylation (A) and Enhancer methylation (B) in ccRCC strongly correlates with enrichment of the EMT pathway, providing a link between SDH loss-induced genome-wide methylation and enhancement of EMT in ccRCC tumors. **C-E**. The Pearson correlation R-value between the log2 expression of *SDHD, SDHB, SDHC* and the log2 expression of E-cadherin (*CDH1*) was 0.56, 0.42, 0.4 respectively (p= 0 for each, KIRC TCGA tumor and normal). **F, G**. Succinate treatment of ccRCC cells (769P) resulted in an increase in the hypermethylated fraction of *CDH1* DNA and under-expression of *CDH1*, both of which were reversed with addition of Ascorbic acid (AA), a cofactor of the TET enzymes (n=2, *: p<0.05, **: p<0.01). **H, I**. In ccRCC cell lines 786-O and 769-P, AA treatment with succinate fully reversed the inhibitory effect of succinate on TET activity (n=2 for each cell line, *: p<0.05, **: p<0.01) TET enzymatic activity was measured by using the ELISA-based Epigenase 5mC Hydroxylase TET Activity/Inhibition Assay Kit (Fluorometric) according to the manufacturer’s instructions. This technique relies on the conversion of methylated products at the bottom of the wells to hydroxymethylated products by the TET enzyme present in the nuclear extract. Thus, the amount of hydroxymethylated products formed is a measure of the TET activity of the nuclear extract harvested from the cells being tested. **J, K**. AA treatment with succinate significantly inhibited succinate-induced invasiveness in ccRCC cell lines 786-O and 769-P (n=2 for each cell line, *:p<0.05; **:p<0.01, 72-hour timepoint). Representative pictures shown. **L**. Knockdown of *TET2* with siRNA resulted in abrogation of succinate-induced increase in invasiveness of ccRCC cells 769P. Knockdown of *TET2*, even without succinate treatment, increased invasiveness in comparison to control. **M**. Western blot showing loss of TET2 in 769P with siRNA knockdown. The complete reversal of succinate –induced TET inhibition, *CDH1* hypermethylation and under-expression, as well as enhanced invasiveness in ccRCC cells by AA, a TET-activating demethylating agent, as well as the abrogation of succinate-induced increase in invasiveness with *TET-2* knockdown, put together, strongly indicate that TET-2 inhibition-induced global regulatory DNA hypermethylation drives SDH loss-induced enrichment of EMT. This was further confirmed by reversal of succinate-induced invasiveness of RCC cells by an archetypal DNA Methyltransferase (DNMT1) inhibitor, azacytidine (Aza; Figure S11). This Aza-induced reversal of invasiveness was associated with an expected reduction in global methylation (Figure S12)

Next, we correlated the expression of subunits *SDHB, SDHC, SDHD* with that of epithelial and mesenchymal markers, and found a very strong positive correlation with E-cadherin (*CDH1*). *CDH1* is an epithelial mark known to be suppressed by promoter methylation in ccRCC, with progressive loss of CDH1 expression and increase in promoter CpG methylation with higher grade in RCC ^11^. The Pearson correlation R-value between the log_2_ expression of *SDHB, SDHC, SDHD* and the log_2_ expression of *CDH1* was 0.42, 0.56, 0.4 respectively (p= 0 for each, Figures 6C-E).

We hypothesized that succinate accumulation from *SDH* loss results in increased *CDH1* methylation and under-expression in ccRCC. Indeed, succinate treatment of ccRCC cells (769-P) resulted in an increase in the hypermethylated fraction of *CDH1* DNA and under-expression of *CDH1*, both of which were reversed with addition of AA, a cofactor of the TET enzymes (Figures 6F, 6G). Similarly, AA co-treatment with succinate of ccRCC cell line 786-O also resulted in enhanced expression of *CDH1* (1.6-fold, n=2, p<0.05, Figure S10), a decrease in total methylated *CDH1* DNA and an increase in total unmethylated DNA, compared to succinate treatment alone.

Next, we aimed to determine whether succinate-induced inhibition of cumulative TET activity in ccRCC cells can be reversed with AA. We found that in ccRCC cell lines 786-O and 769-P, AA co-treatment fully reversed the inhibitory effect of succinate on TET activity (Figures 6H, 6I).

Having shown that succinate promotes global methylation and invasiveness in ccRCC cells, and that AA fully reverses the inhibition of TET activity by succinate, we hypothesized that AA treatment could reverse succinate-induced ccRCC cell invasiveness. Indeed, AA co-treatment with succinate significantly inhibited the succinate-induced invasiveness of both ccRCC cell lines 786-O (p=0.01, n=2, 72-hour time point) and 769-P (p=0.004, n=2, 72-hour time point) (Figures 6J, 6K). Furthermore, this reversal of succinate-induced invasiveness by AA was associated with a dramatic increase in 5hmC levels as shown by dot blot assay (Figure S11).

Knockdown of *TET2* with siRNA resulted in abrogation of succinate-induced increase in invasiveness of 769-P cells. Knockdown of *TET2*, even without succinate treatment, increased invasiveness in comparison to control (Figure 6L).

Therefore, these observations that AA, a TET-activating demethylating agent, caused a complete reversal of succinate–induced TET inhibition, *CDH1* hypermethylation and under-expression, as well as enhanced invasiveness in ccRCC cells, together with abrogation of succinate-induced increase in invasiveness with TET-2 knockdown, put together, strongly indicate that TET-2 inhibition-induced global regulatory DNA hypermethylation drives SDH loss-induced enrichment of EMT. This was further confirmed by reversal of succinate-induced invasiveness of RCC cells by an archetypal DNA Methyltransferase (DNMT1) inhibitor, azacytidine (Aza; Figure S11). This Aza-induced reversal of invasiveness was associated with an expected reduction in global methylation (demonstrated with global DNA Methylation ELISA, Figure S12).

### Fluorescence quenching of recombinant TET-2 protein is unaffected by succinate in the presence of ascorbic acid (AA)

In ccRCC, succinate concentration is higher in tumors compared to adjacent normal renal tissue, whereas fumarate is lower in tumors compared to normal tissue (Figure 2). In a different type of RCC-type 2 Papillary with fumarate hydratase germline mutations (much less common), fumarate accumulation is characteristic. We therefore performed fluorescence quenching experiments with both succinate and fumarate proteins to determine the quenching efficiency of each, as well as their TET-2 protein level interactions with AA.

We first studied the interaction of TET-2 with succinate and fumarate (Figure 7A). The quenching efficiency of fumarate (0.8 +/-0.06 M^-1^) was found to be higher than succinate (0.1 +/-0.005 M^-1^) yielding a p-value of <0.001, suggesting that the substrate specificity of TET2 is higher for fumarate than succinate. Figure 7C shows the comparison of the Stern-Volmer constants ± SEM obtained from the linear range of the data in Figure 6A. In equation y= m × x + c, m indicates the Stern-Volmer constant. The (y, x) points are obtained from Figure 7A.

**Fig 7:**
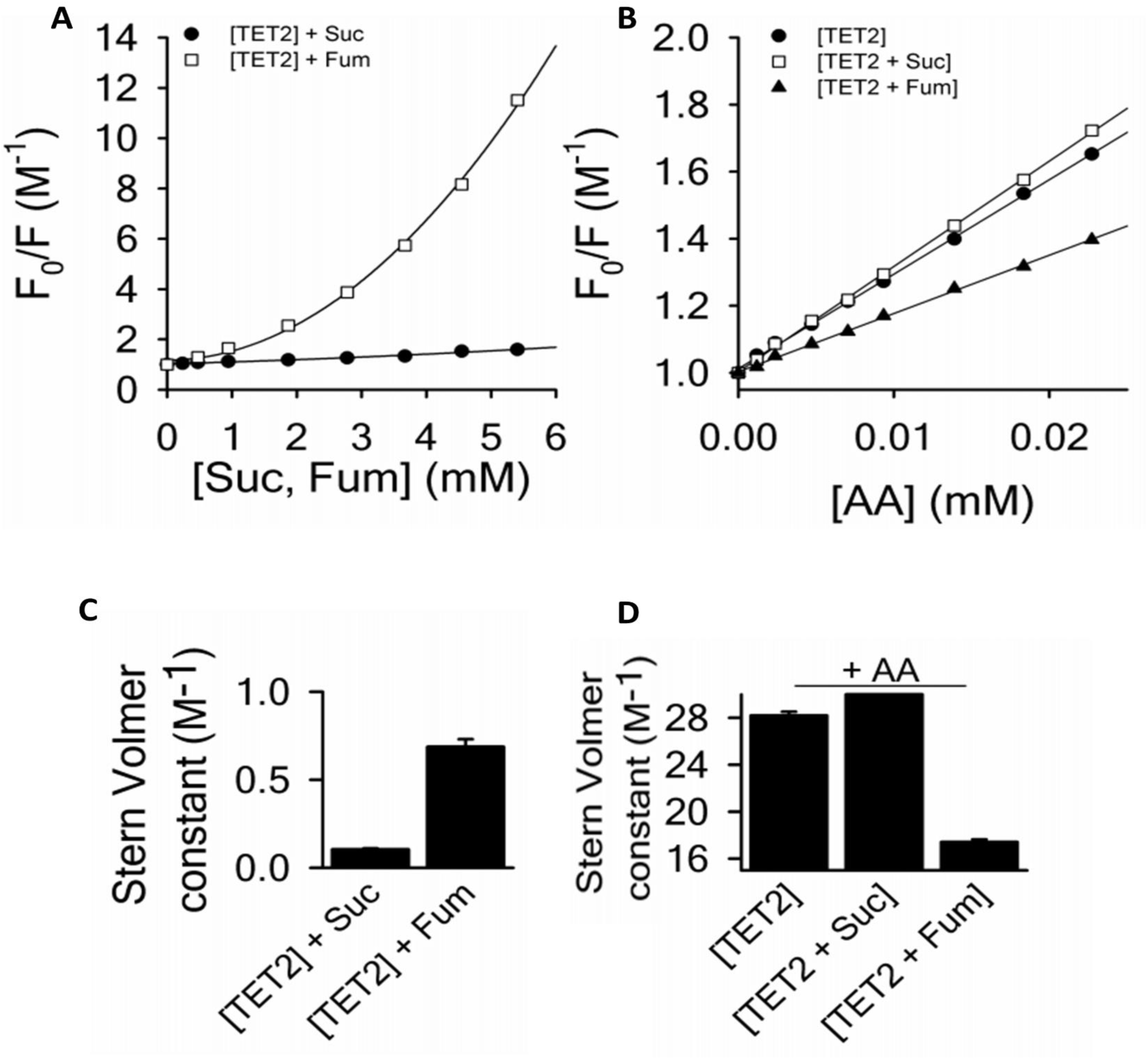
Fluorescence quenching of recombinant TET-2 protein suggests it is unaffected by succinate in the presence of ascorbic acid (AA) **A**. The quenching efficiency of fumarate (0.8 +/-0.06 M-1) was found to be higher than succinate (0.1 +/-0.005 M-1) yielding a p-value of <0.001, suggesting that the substrate specificity of TET2 is higher for fumarate than succinate. **B**. There was an overlap of the quenching curves of TET-2+AA (28.9 +/-0.3 M-1) and TET-2+succinate+AA (31.4 +/-0.2 M-1), suggesting that fluorescence quenching of TET-2 is unaffected by succinate in the presence of AA. However, the quenching curve of TET-2+fumarate+AA (17.4 +/-0.2 M-1) was significantly lower than TET-2+AA (28.9 +/-0.3 M-1) suggesting that fluorescence quenching of TET-2 is affected by fumarate even in the presence of AA. **C**. Comparison of the Stern-Volmer constants ± SEM obtained from the linear range of the data in figure 6A. In the equation y = m × x + c, m indicates the Stern-Volmer constant. The (y, x) points are obtained from figure 6A. **D**. Comparison of the Stern-Volmer constants ± SEM obtained from the linear range of the data in figure 6B. These fluorescence quenching experiments demonstrate the quenching of the fluorescence signal emitted by the TET-2 protein consequent to structural changes induced by the binding of the protein with succinate, fumarate or AA. Put together, these data suggest that in the presence of AA, the TET-2 enzyme may be unaffected by succinate (which is significantly higher in ccRCC compared to adjacent normal kidney, as shown in Figure 2A) but that it may still be affected by fumarate (which is significantly lower in ccRCC compared to adjacent normal kidney, as shown in Figure 2D).

We then performed the quenching of TET-2 with AA, in the presence and absence of succinate and fumarate (Figure 7B). There was an overlap of the quenching curves of TET-2+AA (28.9 +/-0.3 M^-1^) and TET-2+succinate+AA (31.4 +/-0.2 M^-1^), suggesting that fluorescence quenching of TET-2 is unaffected by succinate in the presence of AA. However, the quenching curve of TET-2+fumarate+AA (17.4 +/-0.2 M^-1^) was significantly lower than TET-2+AA alone (28.9 +/-0.3 M^-1^) suggesting that fluorescence quenching of TET-2 is affected by fumarate even in the presence of AA. Figure 7D shows the comparison of the Stern-Volmer constants ± SEM obtained from the linear range of the data in Figure 7B. Put together, these data suggest that in the presence of AA, the TET-2 enzyme may be unaffected by succinate (which is significantly higher in ccRCC compared to adjacent normal kidney, as shown in Figure 2A) but that it may still be affected by fumarate (which is significantly lower in ccRCC compared to adjacent normal kidney, as shown in Figure 2D).

These fluorescence quenching experiments demonstrate the quenching of the fluorescence signal emitted by the TET-2 protein is consequent to structural changes induced by the binding of the protein with succinate, fumarate or AA. The fact that the direct quenching efficiency of succinate on TET-2 is low despite the fact that it significantly inhibits the TET activity (Figures 5I, 5J), suggests that the process of succinate-induced TET inhibition may be more via product inhibition (α-kg dependent dioxygenases including TET enzymes use α-kg as a substrate and convert it to succinate) than via competitive inhibition.

### ccRCC is characterized by an adverse under-expression of ascorbic acid transporter SLC23A1; intravenous ascorbate improves survival in a representative ccRCC xenograft model

Although several aspects of ascorbic acid mechanisms, dosing, administration and delivery have been studied extensively ^12^, expression of ascorbic acid transporters in cancer and its implications has received little consideration. Here, we show that ascorbic acid transporter *SLC23A1* is under-expressed in ccRCC (Figure 8A) and lower expression is associated with significantly worse overall survival (TCGA, HR for high vs low *SLC23A1*= 0.6; log rank p=0.017, Figure 8B). Furthermore, although the expression of the other ascorbate transporter SLC23A2 does not differ significantly from normal renal tissue (Figure 8C), lower expression of *SLC23A2* is also associated with significantly worse overall survival in ccRCC (TCGA, HR for high vs low *SLC23A2*= 0.51; log rank p=0.006, Figure 8D). Interestingly, the logarithmic expression of *SLC23A2* positively correlates with that of *CDH1* (R= 0.18; p<0.001).

**Fig 8:**
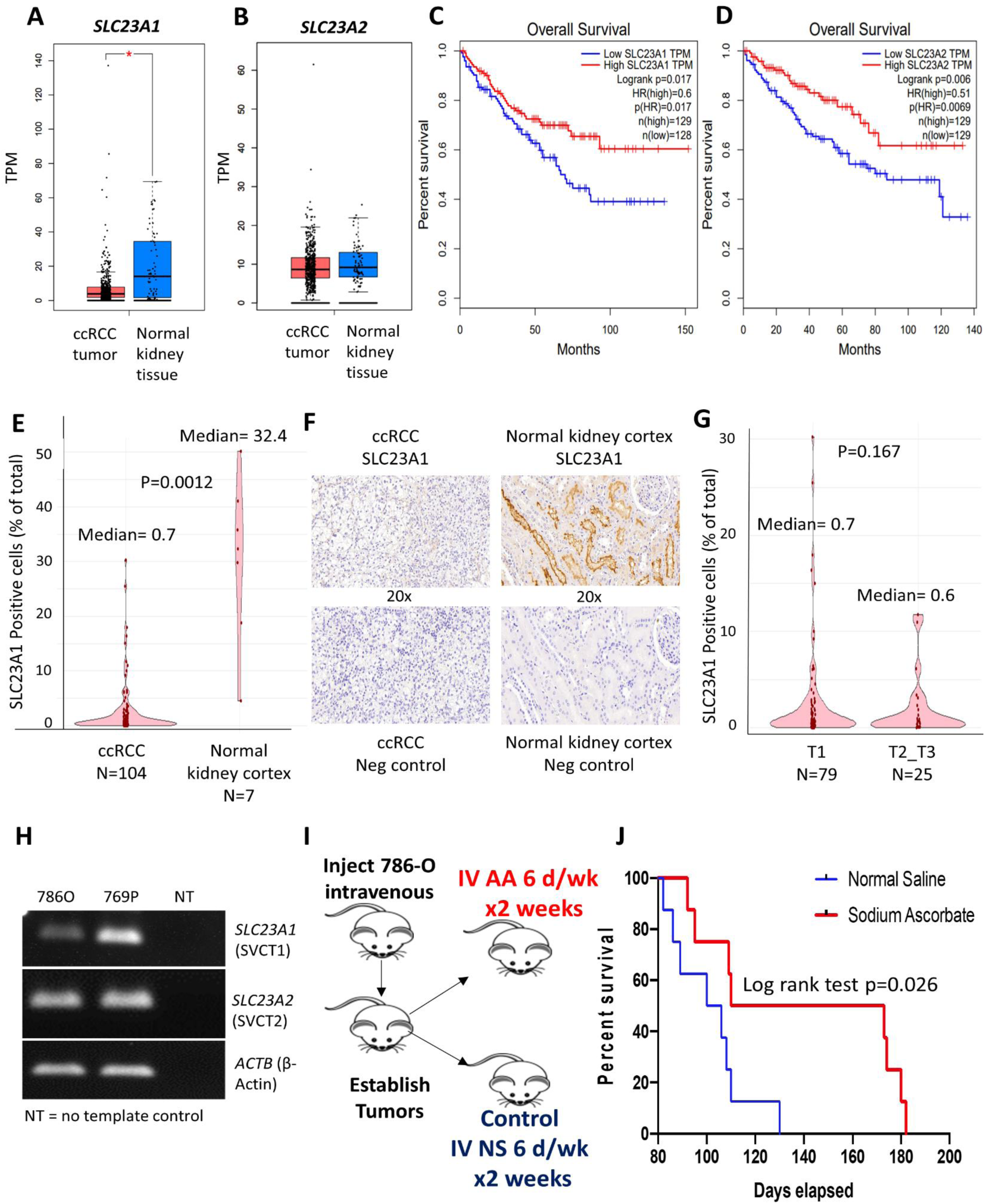
ccRCC is characterized by an adverse under-expression of ascorbic acid transporter SLC23A1; intravenous ascorbate improves survival in a representative ccRCC xenograft model. **A, B**. Ascorbic acid transporter *SLC23A1* (figure 8A), not *SLC23A2* (figure 8B), is under-expressed in ccRCC compared to normal kidney tissue. **C, D**. Lower expression of *SLC23A1* is associated with significantly worse overall survival (TCGA, HR for high vs low SLC23A1= 0.6; log rank p=0.017, figure 8B). Furthermore, although the expression of *SLC23A2* does not differ significantly from normal renal tissue, lower expression of *SLC23A2* is also associated with significantly worse overall survival in ccRCC (TCGA, HR for high vs low SLC23A2= 0.51; log rank p=0.006, figure 8D). **E-G:** Immunohistochemical analysis and artificial intelligence quantitation of SLC23A1 in ccRCC and normal kidney cortex. **E:** ccRCC is characterized by a marked loss of ascorbic acid transporter SLC23A1 [median percent positive cells in ccRCC primary tumors (n=104) and normal kidney cortex (n=7) was 0.7 and 32.4 respectively; p=0.0012]. **F:** Representative IHC images showing high expression in normal kidney cortex tubules and loss of expression in ccRCC. **G:** No significant difference in SLC23A1 expression between T1 (n=79) and T2-T3 (n=25) tumors in ccRCC [median percent positive cells in T1 and T2-T3 tumors was 0.7 and 0.6 respectively; p=0.167]. Taken together, the data indicate that loss of SLC23A1 occurs in the early stages of ccRCC pathogenesis and progression. **H**. PCR products visualized by agarose gel electrophoresis demonstrating reduced *SLC23A1* expression in ccRCC cell line 786-O. **I, J**. We used a metastatic ccRCC xenograft mouse model with human ccRCC cell line 786-O to evaluate the effect of single agent intravenous ascorbate on survival. 786-O has been previously shown to have a higher succinate content than the distal/ subsequent TCA cycle intermediates-fumarate and malate ^13^. Furthermore, 786-O has a markedly low expression of AA transporter *SLC23A1* (Cancer Cell line Encyclopedia and Figure 8H). 3.5 × 10^6^ 786-O ccRCC cells were injected in the tail vein in 16 immunodeficient athymic nude, 6-7 weeks old, male mice. 72 hrs after injection of the ccRCC cells, we randomized the mice into 2 groups of 8 each, one group receiving IV normal saline (6 days/ week for 2 weeks) as control and the other group receiving IV sodium ascorbate (1g/kg 6 days/ week for 2 weeks). At Day 130, there was no mouse alive in the control saline arm, whereas 50% of the mice were alive in the Ascorbate arm. The median survival was 103 and 142 days for the control and ascorbate groups, respectively (log rank test p= 0.026). The percent increased lifespan (%ILS) with IV sodium ascorbate was 38%.

Next, using immunohistochemical analysis and artificial intelligence quantitation of SLC23A1 in ccRCC and normal kidney cortex, we found that ccRCC is characterized by a marked loss of ascorbic acid transporter SLC23A1 [median percent positive cells in ccRCC primary tumors (n=104) and normal kidney cortex (n=7) was 0.7 and 32.4 respectively; p=0.0012, figure 8E, 8F]. There was no significant difference in SLC23A1 expression between T1 (n=79) and T2-T3 (n=25) tumors in ccRCC (figure 8G). Taken together, the data indicate that loss of SLC23A1 occurs in the early stages of ccRCC pathogenesis and progression

We used a representative metastatic ccRCC xenograft mouse model with the human ccRCC cell line 786-O to evaluate the effect of single agent intravenous ascorbate on survival. 786-O has a markedly low expression of *SLC23A1* (Cancer Cell Line Encyclopedia; confirmed with qPCR and visualized by agarose gel electrophoresis-Figure 8H). Furthermore, 786-O has been previously shown to have a higher succinate concentration than distal/ subsequent Krebs cycle intermediates-fumarate and malate ^13^. 3.5 × 10^6^ 786-O ccRCC cells were injected in the tail vein in 16 immunodeficient athymic nude, 6-7 weeks old, male mice. 72 hrs after injection of the ccRCC cells, we randomized the mice into 2 groups of 8 each, one group receiving IV normal saline (6 days/ week for 2 weeks) as control and the other group receiving IV sodium ascorbate (1g/kg 6 days/ week for 2 weeks). At Day 130, there was no mouse alive in the control saline arm, whereas 50% of the mice were alive in the ascorbate arm. The median survival was 103 and 142 days for the control and treatment groups, respectively. The percent increased lifespan (%ILS) was 38%. (log-rank p= 0.026, Figure 8D)

Comparison of Survival Curves of Vc-Na vs. NS

|  |  |
| --- | --- |
| Log-rank (Mantel-Cox) test |  |
| Chi square | 4.928 |
| df | 1 |
| P value | 0.0264 |

## DISCUSSION

Previously, reduced succinate dehydrogenase (SDH) activity resulting in adverse succinate accumulation was thought to be relevant only to 0.05-0.5% of kidney cancers associated with germline *SDH* mutations. In this project, we report that ccRCC (which accounts for ∼80% of all kidney cancers) is characterized by a marked loss of succinate Dehydrogenase subunits *SDHB, SDHC, SDHD* at the mRNA level compared to normal renal tissue, and that this loss is associated with markedly worse overall and disease-free survival in a large cohort of ccRCC patients. We also show, with immunohistochemical staining, that SDHB and SDHD subunits are under-expressed at the protein level in ccRCC compared with adjacent normal renal tubular cells. Using the ccRCC metabolomic repository ^3^, we report that this loss is manifested by the accumulation of succinate during pathogenesis and progression in ccRCC (average 2-fold higher succinate content in ccRCC tumor vs normal; average 1.6-fold higher succinate content in higher stage vs lower stage in ccRCC). The loss of SDH complex prevents the conversion of succinate to fumarate.

Next, we aimed to investigate the mechanism of *SDH* downregulation in ccRCC. We found that Von Hippel Lindau (VHL) loss induced Hypoxia-Inducible factor (HIF)-dependent upregulation of miR210 in ccRCC causes direct inhibition of the *SDHD* transcript. We report that mir-210 is the second-most upregulated miRNA in ccRCC; and of all the miRNAs with a binding site for HIF2α in 786-O, the strength of binding of HIF2α on the mir210 promoter is the highest, by a long margin. Furthermore, higher expression of mir-210 is associated with significantly worse survival in ccRCC (TCGA). Analysis of the TCGA-KIRC dataset revealed that mir-210 expression had a strong negative correlation with *SDH* subunits in ccRCC, which led us to hypothesize that it may be playing a causative role in *SDH* downregulation. Furthermore, *SDHD* is a predicted target of mir-210 (mirDB-microRNA target prediction database). Indeed, mir-210 transfection of immortalized kidney cells (HK2 and HK8) led to marked downregulation of *SDHD* expression (but not of *SDHB* expression), and was found to significantly inhibit *SDHD* 3’UTR activity with a luciferase reporter assay. Put together, these data strongly indicated that mir-210 directly inhibits the *SDHD* transcript in ccRCC pathogenesis. We then hypothesized, and demonstrated, that VHL loss (a pathognomonic feature of ccRCC carcinogenesis) is sufficient to induce *SDH* down-regulation in ccRCC carcinogenesis. CRISPR mediated mono-allelic deletion of VHL in the HK2 cell line increased HIF2α expression, increased mir-210 expression, and decreased *SDHD*/ *SDHB* expression.

Having determined that under-expression of *SDH* subunits is associated with markedly worse prognosis, we then sought to determine the key oncogenic pathway/s enriched with the loss of *SDH*. Analyzing the TCGA-KIRC dataset with the Gene Set Enrichment Analysis (GSEA) software, we found that lower *SDHD* is associated with marked enrichment of the EMT pathway, which aids invasion and metastasis. In fact, EMT was the most enriched pathway in ccRCC tumors with lower *SDHD* expression. We then hypothesized, and demonstrated, that succinate accumulation from loss of SDH aids invasion and metastasis in ccRCC cells, using *in vitro* invasion and migration assays. Using a previously defined EMT score, in which normalized expression values of mesenchymal markers are given a positive score and that of epithelial markers are given a negative score, we found that enhanced EMT portends worse survival in ccRCC.

Next, we sought to investigate the mechanism of EMT enrichment by SDH down-regulation. Succinate has been shown to be an inhibitor of α-kg dependent dioxygenases, including the TET enzymes ^10,14-18^. We therefore hypothesized that loss of SDH and subsequent accumulation of succinate may result in inhibition of TET enzyme activity and global regulatory hypermethylation, which may then play a key role in upregulation of the EMT pathway. We found that loss of *SDHD* and *SDHB* in ccRCC indeed significantly correlates with increase in global genome-wide cytosine methylation (as well as with regulatory region-specific methylation at promoter CpG islands (CGI) and enhancers. Succinate treatment of ccRCC cells resulted in an inhibition of activity of the TET enzymes, explaining the correlation between loss of *SDH* subunits and gain of cytosine methylation in ccRCC. Pathway enrichment analysis revealed that promoter CGI methylation and enhancer methylation in ccRCC strongly correlates with enrichment of the EMT pathway, providing a link between SDH loss-induced genome-wide methylation and enhancement of EMT in ccRCC tumors. We hypothesized that SDH loss-induced regulatory DNA hypermethylation silences ‘epithelial’ marks, such as the E-cadherin gene *CDH1*, thereby favoring the mesenchymal state. *CDH1* is known to be suppressed by promoter methylation in ccRCC, with progressive loss of expression and increase in promoter CpG methylation with higher grade ^11,19,20^. We found that there was a very strong correlation between *SDH* subunits (particularly *SDHD)* and *CDH1* in ccRCC. Succinate treatment of ccRCC cells (769-P) resulted in further increase in the hypermethylated fraction of *CDH1* DNA and under-expression of *CDH1*, both of which were reversed with addition of AA (a TET enzyme-cofactor), explaining the strong correlation between *SDH* subunits and *CDH1*. Furthermore, AA co-treatment fully reversed the inhibitory effect of succinate on TET activity, and also mitigated the effect of succinate on invasiveness in ccRCC cells. This complete reversal of succinate induced-TET inhibition, *CDH1* hypermethylation and under-expression, as well as enhanced invasiveness in ccRCC cells by a TET-activating demethylating agent, strongly indicated that global regulatory DNA hypermethylation drives SDH loss-induced enrichment of EMT in ccRCC. This was further confirmed by reversal of succinate-induced invasiveness of RCC cells by an archetypal DNA Methyltransferase (DNMT1) inhibitor, azacytidine (since AA also has other cofactor functions and there is no commercially available ‘specific’ TET enzyme activator).

The 2016 WHO Classification of Renal Tumors ^21^ identified “succinate dehydrogenase–deficient RCC” as a separate classification, referring mainly to RCC that is associated with germline mutations in any of the SDH subunits. However, our findings reported here strongly indicate that the term “SDH-Deficient” is a misnomer because significant SDH subunit loss (and resulting accumulation of succinate which functions as an oncometabolite) is indeed a common, pathognomonic, feature in ccRCC. We therefore propose that the WHO “SDH-Deficient RCC” entity be renamed as “SDH germline mutation-associated RCC”, to avoid confusion in identifying the small group of RCC patients (0.05-0.5%) with germline mutations in SDH subunits, presenting at a median age of 35 years, and with characteristic pathologic findings of pale-to-eosinophilic cytoplasmic vacuoles and rare-to-occasional cytoplasmic inclusions ^21-26^.

We then studied the dynamics of fluorescence quenching of the recombinant TET-2 protein with succinate (and fumarate), in the presence and absence of ascorbic acid. We found that the quenching efficiency of succinate is lower than that of fumarate, indicating that there is a substrate preference in the TET-2 protein for fumarate than succinate. We then found that the quenching of TET-2 protein with AA was unaffected by the presence of succinate, but was affected by the presence of fumarate. Put together, these findings indicate that in the presence of AA, the TET-2 protein may be unaffected by succinate. Therefore, the fact that the direct quenching efficiency of succinate on TET-2 was found to be low, despite the fact that it significantly inhibits the TET activity, suggests that the process of succinate-induced TET inhibition may be more via product inhibition (α-kg dependent dioxygenases including TET enzymes use α-kg as a substrate and convert it to succinate) than via competitive inhibition.

The evolution of ascorbic acid in cancer treatment with regards to the history, pharmacokinetics, mechanisms of action in relation to the pharmacokinetics and early phase clinical trial data has been reviewed by us.^12^ Although several aspects of ascorbic acid mechanisms, dosing, administration and delivery have been studied extensively, expression of ascorbic acid transporters in cancer and its implications has received little consideration. Here, we show that ascorbic acid transporter *SLC23A1* is under-expressed in ccRCC and lower expression is associated with significantly worse overall survival. Furthermore, although the expression of the other ascorbate transporter *SLC23A2* does not differ significantly from normal renal tissue, lower expression of *SLC23A2* is also associated with significantly worse overall survival in ccRCC. Next, using a metastatic ccRCC xenograft mouse model with a human ccRCC cell line with higher succinate concentration than the distal/ subsequent Krebs cycle intermediates-fumarate and malate ^13^, as well as markedly low expression of AA transporter *SLC23A1*, we demonstrated that high dose intravenous sodium ascorbate significantly improved overall survival. At Day 130, there was no mouse alive in the control saline arm, whereas 50% of the mice were alive in the ascorbate arm. Median survival for control and IV ascorbate groups was 103 days and 142 days respectively. It is possible that the lower expression of transporters may be one of the reasons why intravenous ascorbate is emerging as a promising anti-cancer agent, whereas oral ascorbate had no anti-cancer activity. Thus, further work is warranted on the relationship between transporter expression in cancers and AA dosing requirement for efficacious anti-cancer activity.

In summary, we show that loss of succinate dehydrogenase activity is common in ccRCC, with adverse functional and prognostic implications. Succinate, which accumulates with loss of succinate dehydrogenase, is an important epigenetic modulating oncometabolite in ccRCC, conferring enhanced invasive and migratory ability. Our findings strongly suggest that the “succinate dehydrogenase-deficient renal cell carcinoma” category in the 2016 WHO classification of Renal tumors needs to be renamed “succinate dehydrogenase germline mutation-associated renal cell carcinoma”. We demonstrate herein that a majority of ccRCC are ‘succinate dehydrogenase-deficient’, with resulting accumulation of succinate, which functions as an oncometabolite. Finally, we show that oncogenic effects of succinate in ccRCC can be reversed with pharmacologic AA.

## METHODS

### Cell lines

ccRCC cell lines 786-O and 769-P were purchased from the American Type Culture Collection (ATCC). Cell line authentication was done at ATCC. Cells were cultured in RPMI-1640 media supplemented with 10% v/v Fetal Bovine Serum (FBS) and 1% v/v Penicillin/Streptomycin. HK-2 kidney cells were purchased from ATCC and HKC-8 cells were kindly provided by Lorainne Racusen (Johns Hopkins University, Baltimore, MD). Both cell lines were cultured in DMEM medium supplemented with 10% FBS and 1% Penicillin/Streptomycin.

### Immunohistochemistry and scoring (SDHB and SDHD)

#### Case selection

Paraffin blocks containing clear cell renal cell carcinoma tumor and adjacent non-neoplastic carcinoma were selected for immunohistochemistry from nephrectomy and partial nephrectomy cases resected at Montefiore Medical Center in 2016. All selected cases were diagnosed as clear cell carcinoma, WHO/ISUP grade 2 or 3, confined to the kidney, and had no sarcomatoid or rhabdoid features.

#### Immunohistochemistry

5 micron sections were cut from selected paraffin blocks, mounted on plus slides (Fisher), baked at 60 C for 1 hour, deparaffinized in xylene (2 × 5 minutes), rehydrated in 100% ethanol (2 × 2 minutes) followed by 95% ethanol (2 × 2 minutes), washed twice with deionized water (dH20), treated with 3% hydrogen peroxide for 10 minutes to quench endogenous peroxidase and rinsed twice in dH20. Slides were microwaved in pH 6 antigen retrieval solution (1X Target Retrieval Solution, Agilent Cat # S1699) to bring near to, but not to boiling (approximately 1.5 minutes), placed in a steamer (Oster Model CKSTSTMD5-W) for 30 minutes, cooled at room temperature for 20 minutes, washed twice in dH20, and then in PBS/0.05% Tween 20/1% BSA (PBSTB) for 5 minutes. Slides were wiped dry around but not on the tissue section, which was then surrounded by a hydrophobic barrier made with an ImmeEdge pen (Vector Labs Cat# H-4000). Each section was treated with 4 drops of serum-free protein block (Agilent cat # X0909) for 20 minutes at room temperature, then removed by tapping the slide without washing, followed by incubation with 100 ul primary antibody (succinate dehydrogenase subunit B (SDHB, Abcam #ab14714 diluted 1:2000) or succinate dehydrogenase subunit D (SDHD, Abcam ab203199 diluted 1:200) in antibody diluent (Life Technologies Cat # 003218) with 1% BSA, in a slide box humidified using wet paper towels (Fisher Scientific Cat # 03-448-1 or -2) for 30 minutes at room temperature, blotted with a paper towel, gently rinsed with 2 dips in PBS, washed in PBSTB 1 × 5 minutes, then 2 × 1 minute. 3 drops per slide of secondary antibody (EnVision + labeled polymer-HRP anti-mouse, Dako K4001, or anti-rabbit, Dako K4003) were added and incubated for 30 minutes at room temperature, washed in PBSTB 1 × 5 minutes, then 2 × 1 minute, and stained with diamino benzene (DAB; Cell Marque DAB Substrate Kit Cat # 957D-60) for 4 minutes. Slides were then washed 4 times in tap water, counterstained with Harris hematoxylin (Leica cat # 3801561), blued with 2 dips in an ammonia solution (1:1000 dilution of stock ammonium hydroxide, (Fisher cat# A669S-500), washed once more in tap water, dehydrated through ethanol and xylene and coverslipped with Cytoseal XYL (Thermo Scientific cat# 8312-4).

#### Evaluation of immunostaining

Staining was evaluated semi-quantitatively on a scale of 0 to 4+. Groups of non-neoplastic tubule or carcinoma cells within contiguous areas were evaluated together, since there was relatively little variation in staining (less than 1+ in the 0 to 4+ scale) among them (supplemental figure). Digital photography was performed with a Nikon Digisite DS-Fi3 microscope camera at a resolution of 2880 × 2048 pixels per image, using the same exposure and gain (contrast) settings for all slides.

### Immunohistochemistry and scoring (SLC23A1)

Immunohistochemistry was performed by Reveal Biosciences on FFPE Kidney Cancer tissue microarrays (TMAs; KC01, KC02, KC04, Reveal Biosciences). TMAs were stained using a Leica Bond automated immunostainer and a rabbit anti-SLC23A1 antibody (PA5-61555, Thermo Fisher) at a dilution of 1:800 plus a negative, no primary antibody control. Heat induced antigen retrieval was performed using Leica Bond Epitope Retrieval Buffer 1 (Citrate solution, pH 6.0) for 20 minutes. Endogenous peroxidase was blocked using Novolink Peroxide block (Leica) for 10 minutes. Non-specific antibody binding was blocked using Novolink Protein block for 20 minutes. Primary antibody was applied for 30 minutes. Anti-rabbit Poly-HRP-IgG secondary antibody (Leica, cat#RE7280-CE_ was applied for 20 minutes. Anti-SLC23A1 antibody was detected using Novocastra Bond Refine Polymer Detection and visualized with 3’3-diaminobenzidine (DAB; brown). A Hematoxylin nuclear counterstain (blue) was applied. Whole slide images were visualized and percent positive staining was quantified for each core using ImageDx software (Reveal Biosciences).

### CRISPR/Cas9 *VHL* knockdown

Three synthetic single guide RNAs (sgRNAs) were designed targeting the first exon of VHL corresponding to the following nucleotide sequences: CGCGGAGGGAATGCCCCGGA; TGAAGAAGACGGCGGGGAGG; and GGAGGAACTGGGCGCCGAGG using the Synthego design tool (Synthego). Cas9 enzyme and sgRNAs were co-transfected using Lipofectamine CRISPRMAX (Invitrogen) following manufacturer’s protocol (Synthego). Monoclonal cell populations were generated from the resulting transfection using limiting dilution and screened for knockdown efficiency by VHL western blot and Sanger sequencing.

### miRNA transfection

Pre-miRNA precursor molecules for *hsa-miR-210-3p* and negative control 1 (Ambion/Thermo Fisher Scientific) were transfected into HK2 and HKC8 cells at a final concentration of 30 nM using Lipofectamine 3000 (Invitrogen). All transfections were performed in triplicate and validated by qPCR for hsa-miR-210-3p.

### SDH Activity Assay

At 48 h post-transfection with miRNA, cells were harvested and assayed immediately for SDH activity. SDH activity of transfected HK2 and HKC8 cells was determined using the Succinate Dehydrogenase Activity Colorimetric Assay Kit following manufacturer’s instructions. Reaction was started by adding a blue colored artificial probe to accept electrons from the succinate to fumarate oxidation. Absorbance at 600 nm were recorded in kinetic mode at room temperature. The decrease in absorbance per unit time (slope) was compared to a standard curve obtained with known protein concentrations and reported proportional to SDH activity using the equation: 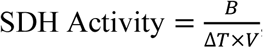, where *B* is the nmol of protein reduced corresponding to the change in absorbance Δ*OD* in the standard curve.

### Immunoblotting

Proteins were isolated from cell lysates and run on 10% Tris-HCl poly-acrylamide gels. Proteins were transferred to nitrocellulose membranes and blocked with 5% non-fat dry milk/TBS containing Tween-20. Membranes were incubated in the following primary antibodies: HIF2α (Novus Biologicals NB100-122), VHL (Novus Biologicals NB100-485), TET2 (Cell Signaling Technology, 18950) and β-Actin (Novus Biologicals, NB600-501; AC-15). Proteins of interest were detected using IRDye 800CW and 680RD secondary antibodies (LI-COR) on a LI-COR Odyssey Fc Imaging System.

### 3-UTR Reporter Assay

*SDHD* 3’UTR LightSwitch Luciferase Reporter Assay was purchased from Active Motif. Cells were co-transfected with 30 nM miRNA and 100 ng reporter using Lipofectamine 3000 in a 96-well plate. Luciferase activity was measured using the LightSwitch Luciferase Assay Reagent (Active Motif) on a SpectraMax M5 plate reader (Molecular Devices) following manufacturer’s instructions.

### ChIP-seq Analysis

ChIP-seq data were downloaded from GSE34871^4^, GSE86092^5^, and GSE67237^27^. ChIP-seq data were visualized in IGV^28^. For analysis of HIF2α and HIF1α binding site distribution, ChIP-seq files were analyzed using *ChIPSeeker* ^29^.

### RNA-seq analysis

RNA sequencing data were accessed from GSE1078484^6^ using GEO RNA-Seq Experiments Interactive Navigator (GREIN ^30^).

### Bioinformatic analyses

The <u>Gene Expression Profiling Interactive Analysis (GEPIA)</u> ^31^ was used for the following: *SDHA, SDHB, SDHC, SDHD, FH, MD, CDH1, SLC23A1, SLC23A2* expression comparison between ccRCC tumors and normal kidney (TCGA-KIRC dataset, Figures 1,2,5); Survival analyses with *SDHA, SDHB, SDHC, SDHD, CDH, SLC23A1, SLC23A2* expression (TCGA-KIRC dataset, Figures 1,5); correlation between *SDHB, SDHD, SDHC* expression with *CDH1* in ccRCC tumors and normal kidney (TCGA-KIRC dataset, Figures 1,5).

The <u>TCGA Wanderer</u> platform ^32^ was used for the following: Promoter CGI methylation of *SDHB, SDHC, SDHD* comparison between ccRCC tumor and normal (TCGA-KIRC dataset, Figure S3); correlation between *CDH1* expression and methylation beta value at CGI probe cg17655614 in ccRCC (TCGA-KIRC, Figure 5).

The Gene Set Enrichment Analysis (GSEA) platform ^33,34^ was used for Pathway Enrichment Analysis. ccRCC KIRC data was downloaded from TCGA. A ranked dataset comparing two groups (tumor vs normal, Fig S4; or SDHB high vs SDHB low, Fig 4) was generated by calculating the product of the fold-change sign and –log(P-value) for each gene. This ranked file was uploaded in the GSEA platform for analysis of pathways that were for these comparisons

### Invasion Assay

BD BioCoat Matrigel Invasion Chambers were used to assess the invasiveness of tumor cells, with control, succinate alone (50uM) or together with Ascorbic acid (1mM) or Azacytidine (1uM). A thin layer of Matrigel matrix at the bottom of each chamber serves as a reconstituted basement membrane *in vitro*, while the chemoattractant is present in the culture medium on the outside of the chamber. Manufacturer’s protocol to prepare the Matrigel chambers was followed and same number of cells per chamber (at least 20,000 cells) were plated in replicates. Incubation time of the cells in the Matrigel chambers was 72 h after which the bottom of the chambers was fixed with buffered formaldehyde, stained with bromophenol blue. The number of invading cells were counted after removal of Matrigel layer and reported as mean +/-SEM.

### Scratch Assay

Cells were grown to ∼60% confluence under each treatment condition (succinate and control) in 6 well plates. After 18 h treatment, scratches were made using a 200 μl pipet tip, at least 5 mm apart on the cellular monolayer in each well. This was marked as t = 0 h point. The wells were quickly washed with PBS and replenished by fresh medium. Bright field images were captured at the indicated times as cells migrated towards the scratched area to close the wound. A fixed-dimensions rectangular field was demarcated using imaging software ImageJ such that the field contained no cell at t = 0 h. The number of cells migrating into the field were counted at each timepoint (with a blue dot over each migrated cell as shown in figure 4D) and reported.

### EMT score

To avoid bias in the selection of markers, we used a previously defined EMT score (6), in which normalized expression values of mesenchymal markers (FN1, VIM, ZEB1, ZEB2, TWIST1, TWIST2, SNAI1, SNA2, CDH2) are given a positive score and that of epithelial markers (CLDN4, CLDN7, TJP3, MUC1, CDH1) are given a negative score. The TCGA-KIRC dataset was analyzed for these genes and the composite EMT score divided (based on median) into ‘high’ and ‘low’ EMT scores. These two groups were then compared for overall survival.

### Nuclear protein extraction and *in vitro* TET enzymatic activity analysis

Cells were treated with different conditions for 24 h prior to harvesting by trypsinization. Nuclear protein was then isolated from cells using the EpiQuik Nuclear Extraction Kit (EpiGentek Group Inc.), according to the manufacturer’s instructions. TET enzymatic activity was measured by using the ELISA-based Epigenase 5mC Hydroxylase TET Activity/Inhibition Assay Kit (Fluorometric) according to the manufacturer’s instructions. This technique relies on the conversion of methylated products at the bottom of the wells to hydroxymethylated products by the TET enzyme present in the nuclear extract. Thus, the amount of hydroxymethylated products formed is a measure of the TET activity of the nuclear extract harvested from the cells being tested. Incubation time of nuclear lysates was 90 minutes. Eight microliters of nuclear lysate (total nuclear protein concentration in excess of 2.5 µg/µl) was used per well for measurement of TET activity.

### Methylation PCR assay

Genomic DNA was extracted from ccRCC cells using the DNeasy Blood and Tissue Kit (Qiagen). Epitect Methyl II PCR Array System (Qiagen) was used to determine the CpG island methylation status of *CDH1* (E-Cadherin). At least 105 ng of total genomic DNA was used per sample. DNA was equally distributed among the parallel digests containing the mock, methylation-sensitive, methylation-dependent and total digestion.

Predesigned primers for *CDH1* locus were obtained along with the kit and quantitative PCR applied to each of the digests. Data analysis was carried out using the manufacturer’s protocol (analogous to ΔΔ_Ct_ method in RT-PCR) which provide gene methylation extent of the total input gDNA as percentages of unmethylated (UM), methylated (M), and hyper-methylated fraction of input DNA.

### Fluorescence spectroscopy

All fluorescence measurements were performed at 20 °C on a Horiba Jobin-Yvon Fluorolog 3 spectrofluorometer equipped with a Wavelength electronics Model LFI-3751 temperature controller. Protein fluorescence emission spectra of TET2 were averaged three times between 305 and 400 nm with excitation at 280 nm. The step width was 1 nm and the integration time 1 s.

All protein solutions contained 0.5µM TET2 in PBS buffer. PBS-buffered stock solutions of succinate, fumarate or ascorbic acid were added stepwise to the protein solutions prior to the measurement of the fluorescence spectra.

Quenching constants were obtained using the Stern-Volmer equation ^35^.

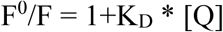

In this equation, K_D_ is the quenching constant and [Q] is a defined concentration of ascorbic acid. F^0^ and F are the fluorescence intensities (at 328 nm) in absence and presence of ascorbic acid respectively.

### Quantitative real time PCR

*<u>Figures 3F, 3K, S4B</u>*: At 48 h post-transfection with miRNA, cells were harvested for RNA extraction. Total RNA was isolated using the miRNeasy kit (Qiagen) following manufacturer’s instructions. cDNA was synthesized using SuperScript III (Invitrogen) and Taqman MicroRNA Reverse Transcription Kit (Thermo Fisher) for mRNA and miRNA analyses, respectively. Taqman primer probe assays (Thermo Fisher) were used for RNU6B (001093), hsa-miR-201-3p, SDHD (Hs00829723_g1), and SDHB (Hs00268117_m1). Quantitative PCR was performed for RNU6B, hsa-miR-201-3p, SDHD, and SDHB using TaqMan Universal Master Mix II (Thermo Fisher), following manufacturer’s instructions. Relative expression of each gene was determined using the ΔΔCt method where *RNU6B* and *GAPDH* were used as housekeepers for miRNA and mRNA genes, respectively.

*<u>Figures 7I, S10A</u>*: Total RNA was isolated from ccRCC cells (786-O and 769-P) using RNeasy Mini Kit (Qiagen), followed by first-strand cDNA synthesis from 2 to 2.5 μg of RNA using SuperScript VILO (Invitrogen). Gene expression of *CDH1* was measured via RT-PCR using forward (5’-GAA CAG CAC GTA CAC AGC CCT-3’) and reverse (5’-GCA GAA GTG TCC CTG TTC CAG-3’) primers. The expression values were normalized to the geometric mean of *GAPDH* (Forward primer: 5’-ACC CCT GGC CAA GGT CAT CCA-3’, Reverse primer: 5’-ACA GTT TCC CGG AGG GGC CA-3’).

*<u>Figure</u> <u>S12</u>*: cDNA was reverse transcribed from RNA using SuperScript III (Invitrogen). PCR was performed using HotStarTaq (Qiagen). Primers sequences for human *SLC23A1* (*SVCT1*), and *ACTB* (β–Actin) were as previously published ^36^. Primers used for human *SLC23A2* (*SVCT2*), Forward/Reverse: CAGGCCAGTGCTTTTGCATT/ TCCGGGGATACCAGATGTGT.

### Dot blot assay for 5-hydroxymethylcytosine analysis

Genomic DNA from the indicated cell lines and treatment conditions (72 hour time point) was extracted. DNA was diluted in total 8 µl water, denatured by adding 2 µl of 2 N NaOH/50 mM EDTA, and 10 minutes incubation at 95 °C. Samples were quickly moved to ice followed by neutralization using 10 µl of ice cold 2 M ammonium acetate. Denatured DNA was manually spotted (769P-200ng; 786O-200ng) on a positively charged nylon membrane (Amersham Hybond N+). Hybridization was carried out by baking for 30 minutes at 80 °C. The membranes were blocked in 5 % milk/TBST for 1 h followed by primary antibody blotting at 4 °C overnight incubation – 1: 2000 rabbit anti-hmC (cat# 39069, Active Motif). The next day the blots were washed with TBST followed by incubation with secondary 1:2000 anti-rabbit-HRP for 1 h, washed, and imaged using Western chemiluminescent reagents on a photographic X-ray film. Methylene blue stain (0.04% in 0.5 M sodium acetate, pH 5.2) was used as loading control. Photographic films and membrane with methylene blue spots were scanned as 8-bit black and white TIFF images on a scanner. Each spot was quantified using ImageJ by subtracting local background from the intensity readout from a fixed-size circle drawn around each spot.

### Quantitative global 5-methycytosine assay (ELISA)

Genomic DNA was used to measure percent cytosine modification to 5-methylcytosine using a one-step ELISA method (cat# 1030, Epigentek). Manufacturer’s protocol was followed to quantify the amount of modified cytosine residues in 50 ng of genomic DNA per treatment condition. Briefly, the protocol involves binding of fixed amount of DNA to each well of a 96-well plate, followed by probing with the supplied anti-5mC antibody and colorimetric measurement at 450 nm representative of the concentration of modified DNA in the well. Readouts in duplicates were reported with standard error of the mean.

### *In vitro* Ascorbic acid (AA) treatment

Immediately prior to *in vitro* treatment with high dose L-ascorbic acid (Sigma-Aldrich, Cat# 50-81-7), cells were exposed to catalase to quench free radicals as previously published ^9,37^. 100 μg/ml catalase (Sigma) in 50mM potassium phosphate was applied to all cells prior to treatment with or without 1 mM ascorbic acid (Sigma). Both catalase and ascorbic acid were prepared fresh for each experiment.

### *In vivo* studies with AA

3.5 × 10^6^ ccRCC cells (786-O) were injected into the tail vein of 16 immunodeficient athymic nude, 6-7 weeks old, male mice, obtained from Jackson Labs. The mice were randomized into two groups of 8 each-‘AA’ group and ‘control’ group. 72 hours after injection of ccRCC cells, treatment was initiated with tail vein injections: IV AA at 1g/kg or IV normal saline (6 days/ week for 2 weeks). Following the 2-week treatment period, mice were monitored on a daily basis until end of study.

### Statistical analyses for clinical data, *in vitro* experiments and *in vivo* xenograft study

#### Clinical data

Paired t-test was used for statistical analyses in figure 1B (SDHB, SDHD IHC intensity scoring comparison between ccRCC and adjacent normal). Log-rank test was used for survival analyses with the Gene Expression Profiling Interactive Analysis (GEPIA) platform. One-sided Student’s T test was used for statistical analyses in figure S3 (promoter CGI methylation comparison)

#### *In vitro* experiments

One-sided Student’s t test was used for statistical analyses in figures 3,4,6,7.

#### *In vivo* xenograft study

Log-rank test was used for statistical analysis of survival difference. Statistical significance was considered P < 0.05.

### Study approvals

The use of ccRCC primary tumors and adjacent normal renal tissue for immunohistochemistry and the ccRCC xenograft study were approved by the Albert Einstein College of Medicine Institutional Review Board (Title: “Molecular Profile Analysis of the Human Kidney”-PI: Shenoy; Lead Pathologist: Pullman) and IACUC, AECOM (PI: Zou).

## AUTHOR CONTRIBUTIONS

Conceptualization and direction: N.S

Data acquisition: R.A, Y.Z, R.A.L, N.S, K.P, N.A, N.R, J.M.A, J.P, J.Y, X.W, L.L, V.M, A.T, S.A, S.G, A.A

Data analysis and interpretation: N.S., R.A., R.L., Y.Z., J.P., K.P., V.M., A.T. IHC and scoring (clinical cases): J.P, J.M.A

Bioinformatics: N.S., K.P.

Fluorescence quenching: V.M, A.T, N.S Xenograft study: Y.Z

Manuscript preparation: N.S

Manuscript editing/ review: N.S, S.H, J.P, R.A.L, B.A.G

## FUNDING

Albert Einstein College of Medicine start-up grant (N.S.); 2017 American Society of Hematology Research Award (N.S.)

## ACKNOWLEDGEMENTS

1. The ccRCC metabolomic repository generated by MSKCC ^3^ and made publicly available via supplemental attachments. The metabolite data in Figure 2 of this manuscript was derived from analysis of this repository.
2. The Cancer Genome Atlas (TCGA) (https://www.cancer.gov/tcga)^38^
3. Interactive Web-based platforms for analysis of the TCGA database: GEPIA (Gene Expression Profiling Interactive Analysis)^31^, TCGA Wanderer ^32^
4. GSEA pathway analysis software

## FIGURE LEGENDS

**Fig S1.**
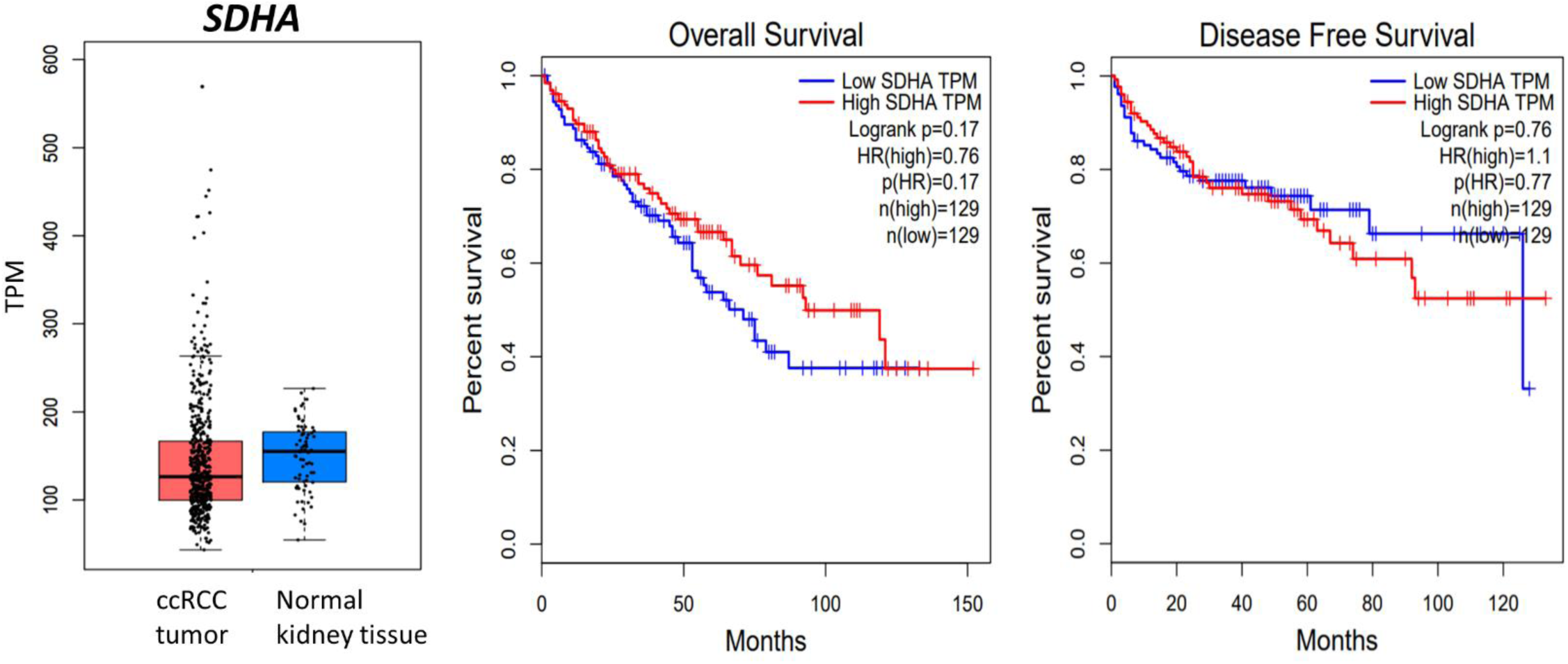
*SDHA* expression in ccRCC is not different from normal kidney tissue (TCGA) and does not impact survival.

**Fig S2.**
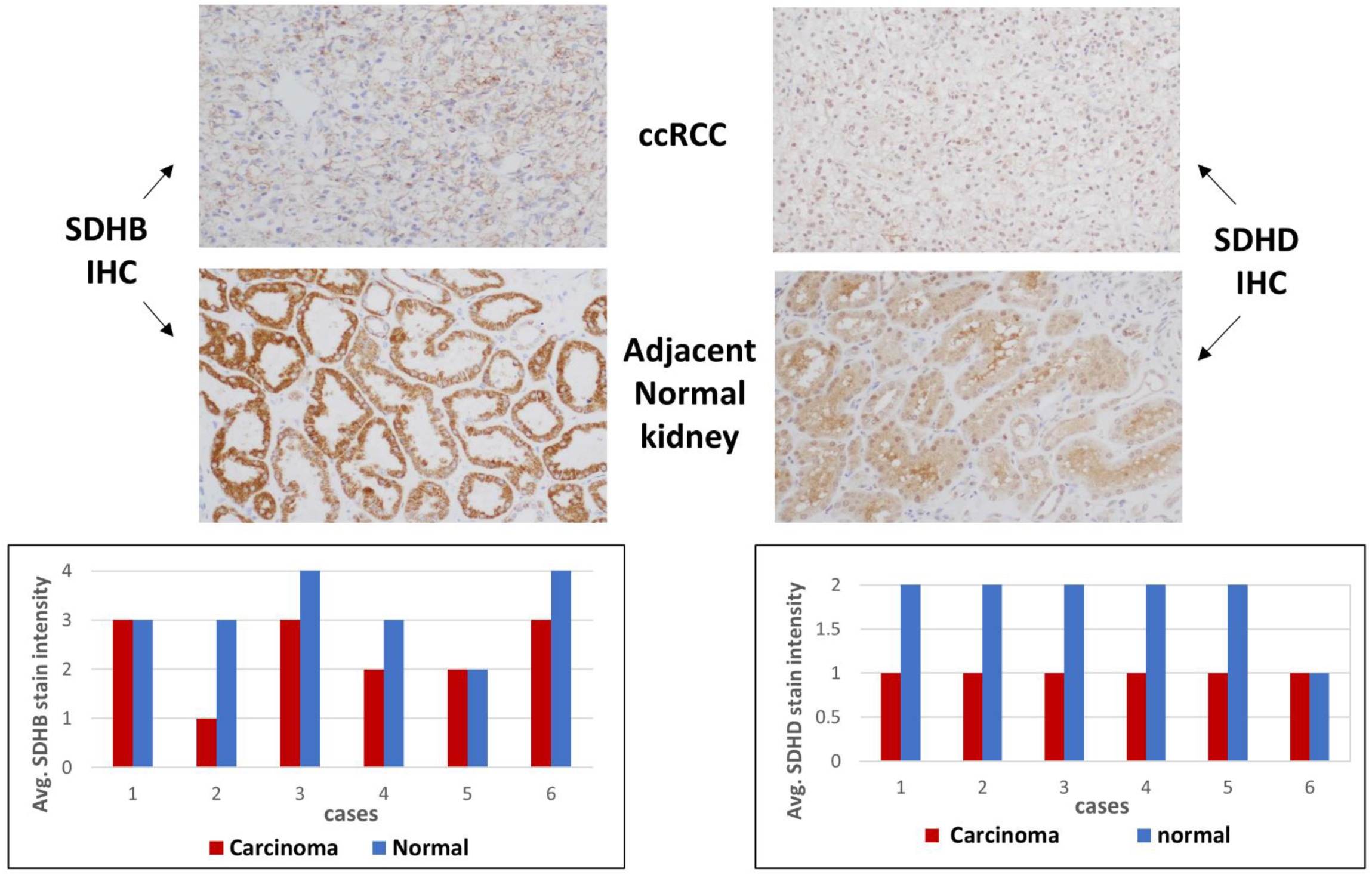
SDHB and SDHD IHC in ccRCC tumors and adjacent normal kidney tissue. Original objective magnification 20X. Bar graphs compare average stain intensities of ccRCC vs adjacent normal renal tubules for 6 different cases. Average intensities of all 6 cases represented in Figure 1D,E graphs. Case 2 illustrated here.

**Fig S3.**
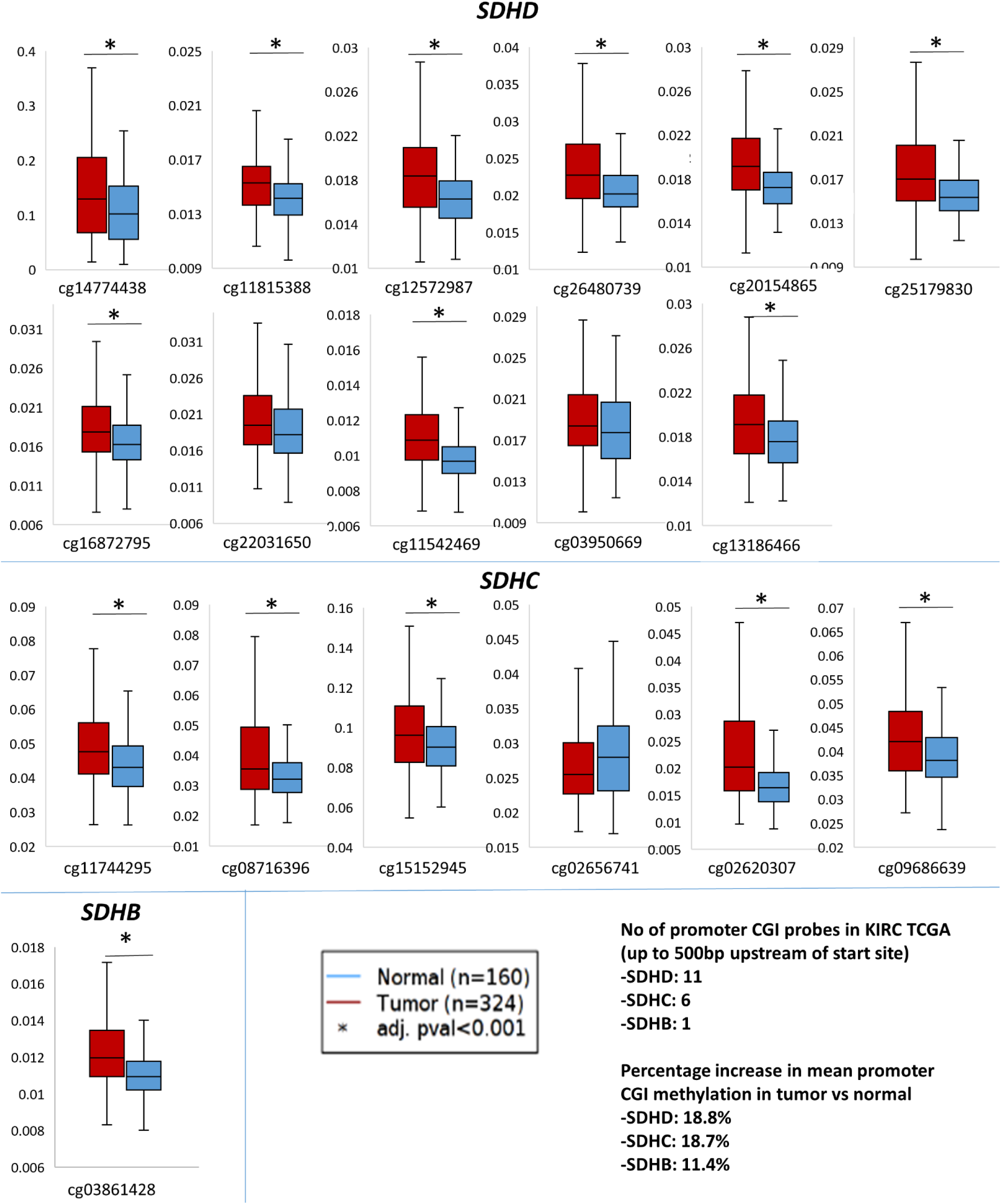
SDH subunits (B,C,D) under-expression is associated with increase in promoter CpG island (CGI) methylation in ccRCC.

**Fig S4.**
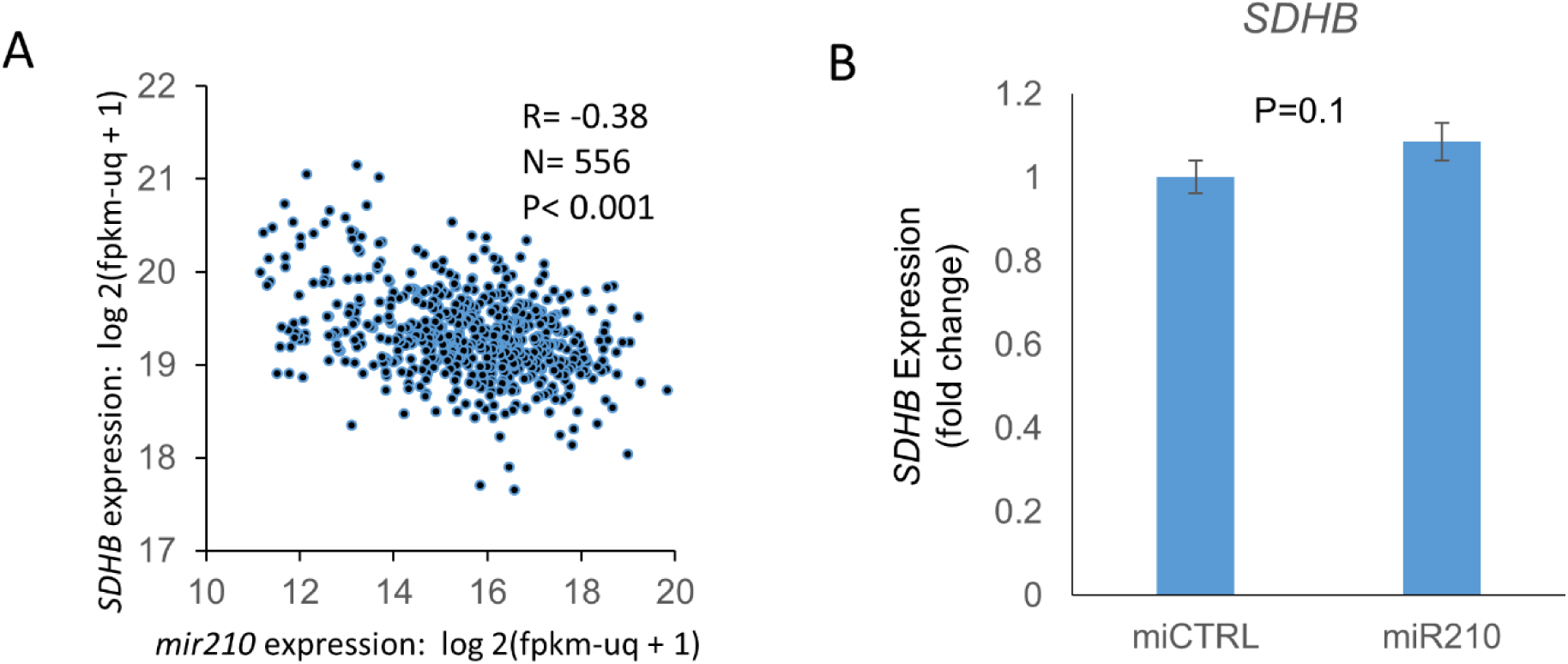
A. Inverse correlation between *mir210* expression and *SDHB* expression (R=-0.38, TCGA). B. miR210 or control miRNA were transfected into HKC8 normal kidney cells. *SDHB* gene expression was measured 48 hours post-transfection. miR210 did not inhibit *SDHB* expression in the HKC8 cell line.

**Fig S5.**
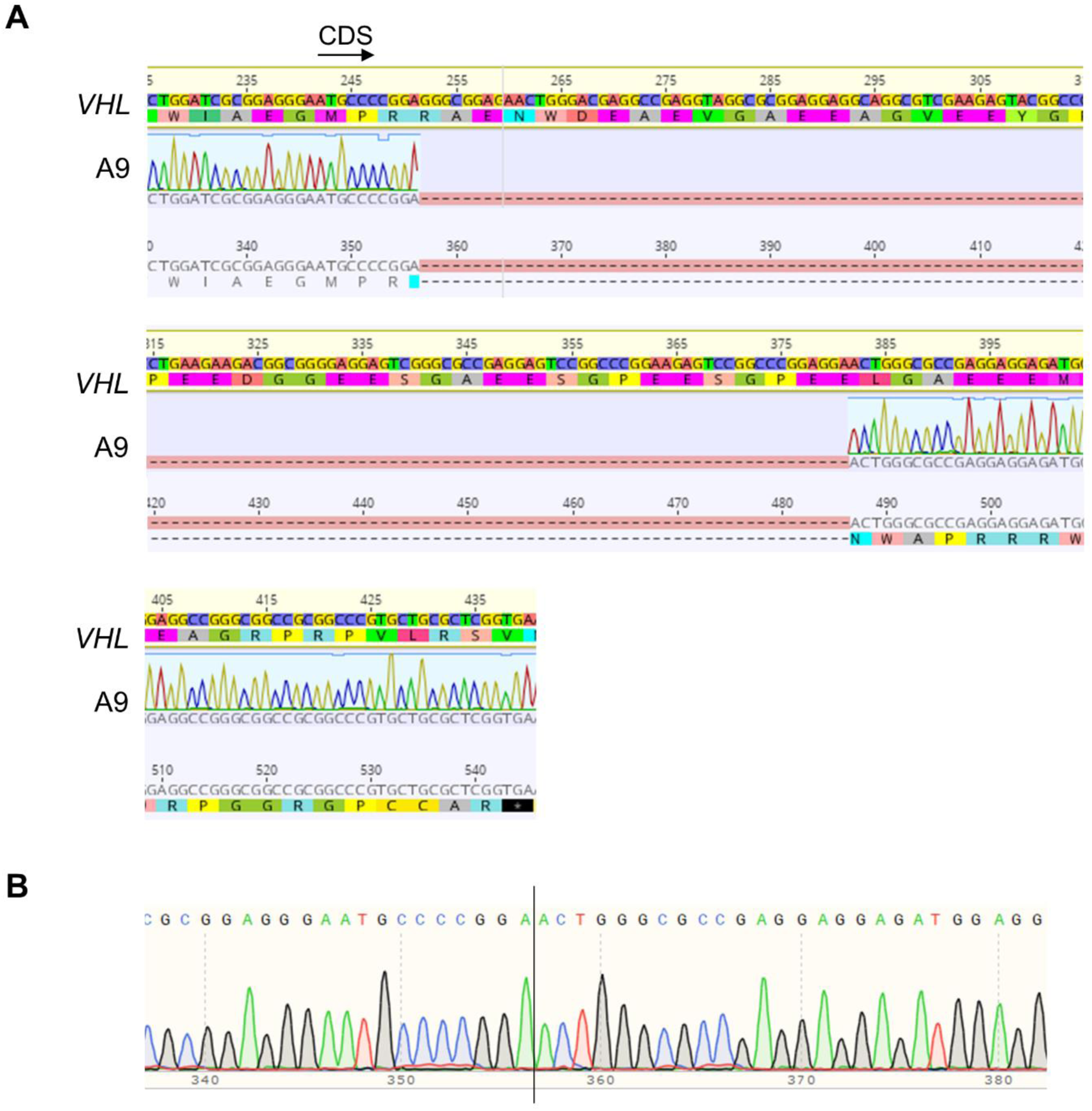
Sanger sequencing confirmation of CRISPR induced mono-allelic *VHL* deletion in A9 clone. A. Alignment to *VHL* genomic sequence shows 130 bp deletion in Exon 1 resulting in a frame-shift and introduction of an early stop codon. B. Sanger sequence of deleted region.

**Fig S6.**
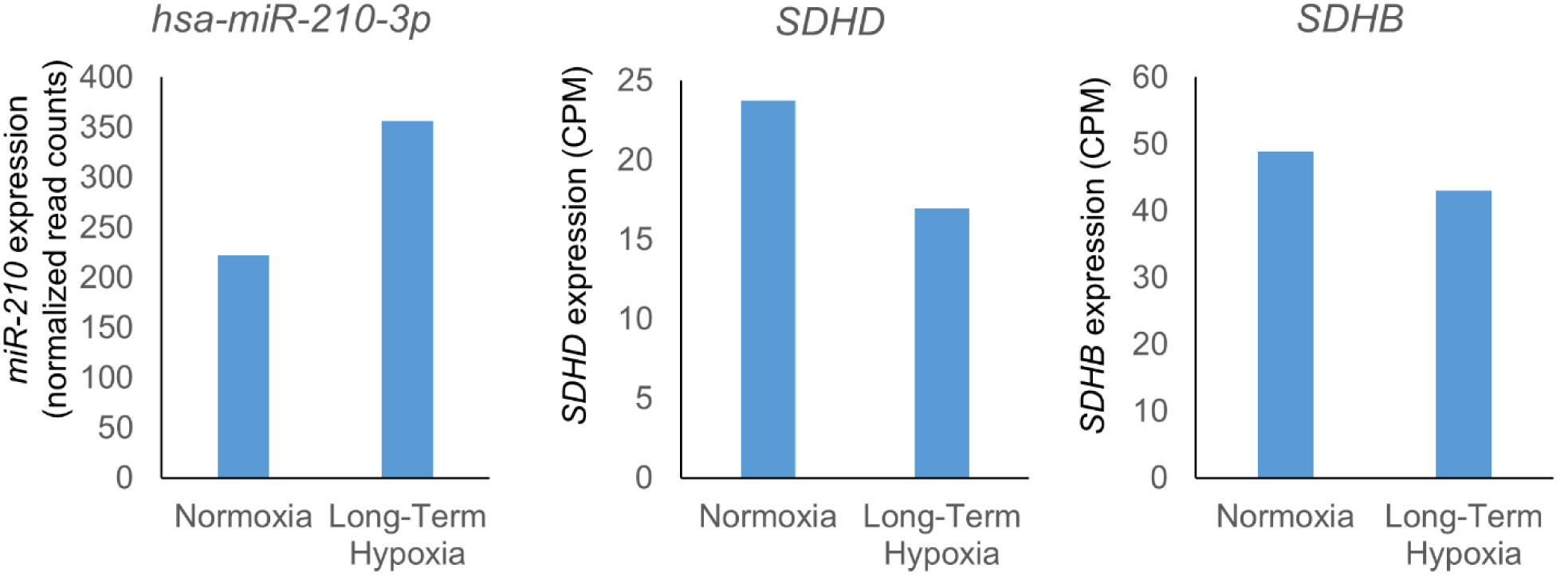
7860 cells were cultured under normoxic (20% 0_2_) or long term hypoxic (1% 0_2_, 3 months) conditions. miRNA and mRNA sequencing were performed for each condition. Data were accessed through GSE107848 ^6^

**Fig S7.**
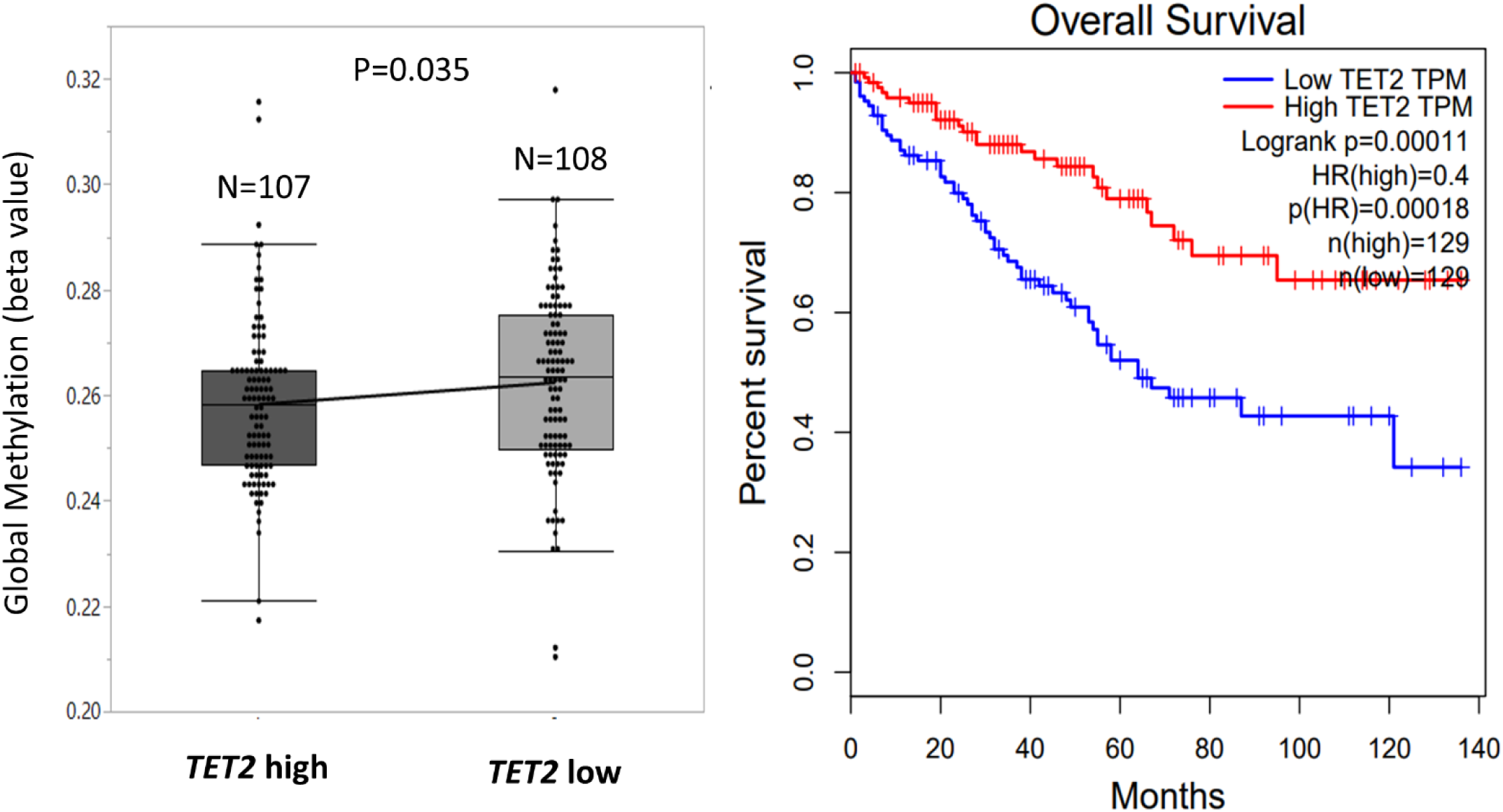
Low TET2 expression is associated with increase in cytosine methylation and worse survival in ccRCC.

**Fig S8.**
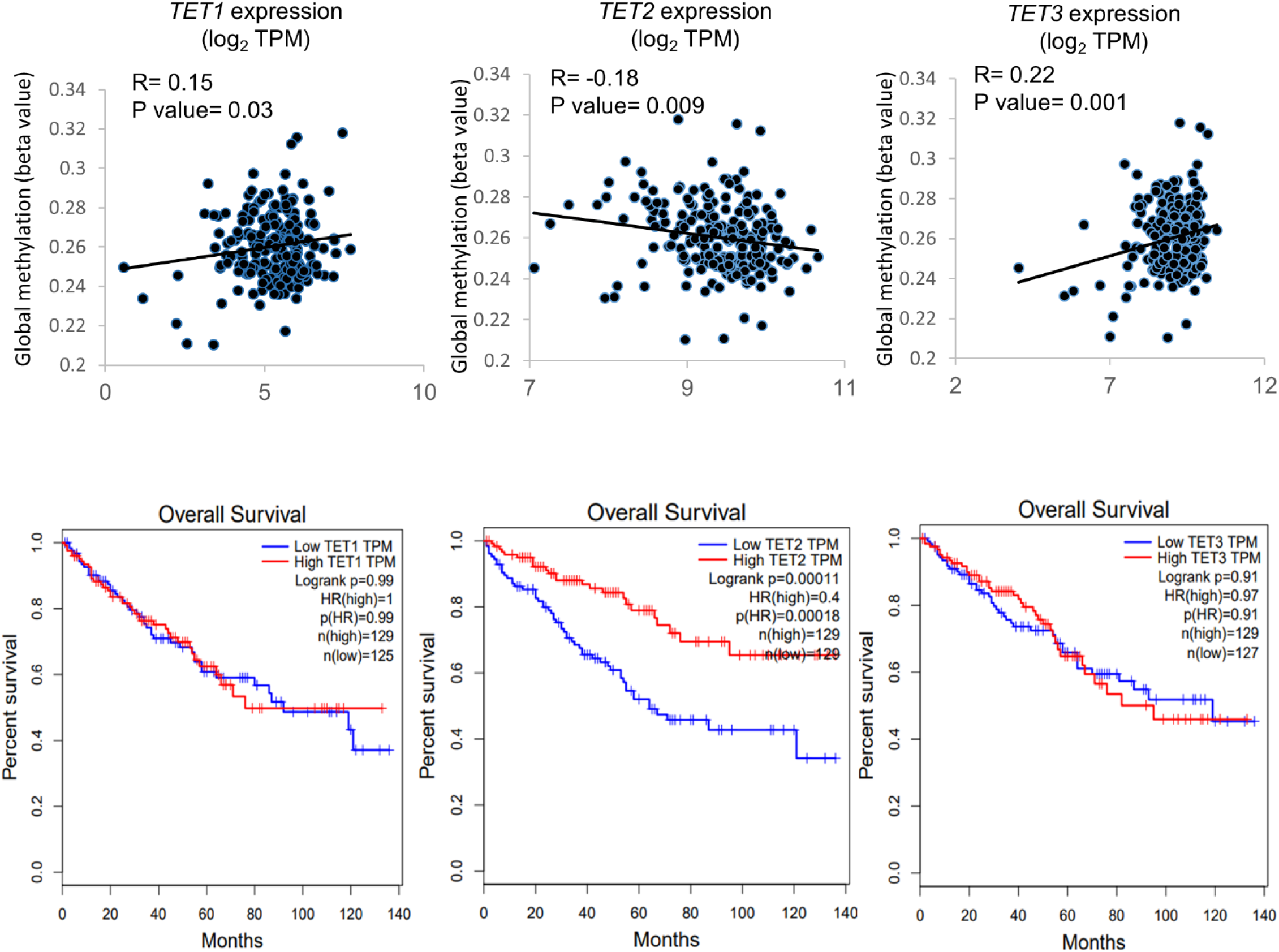
Only *TET2*, not *TET1* or *TET3*, is negatively correlated with global methylation in ccRCC as well as significantly associated with adverse outcome (TCGA)

**Fig S9.**
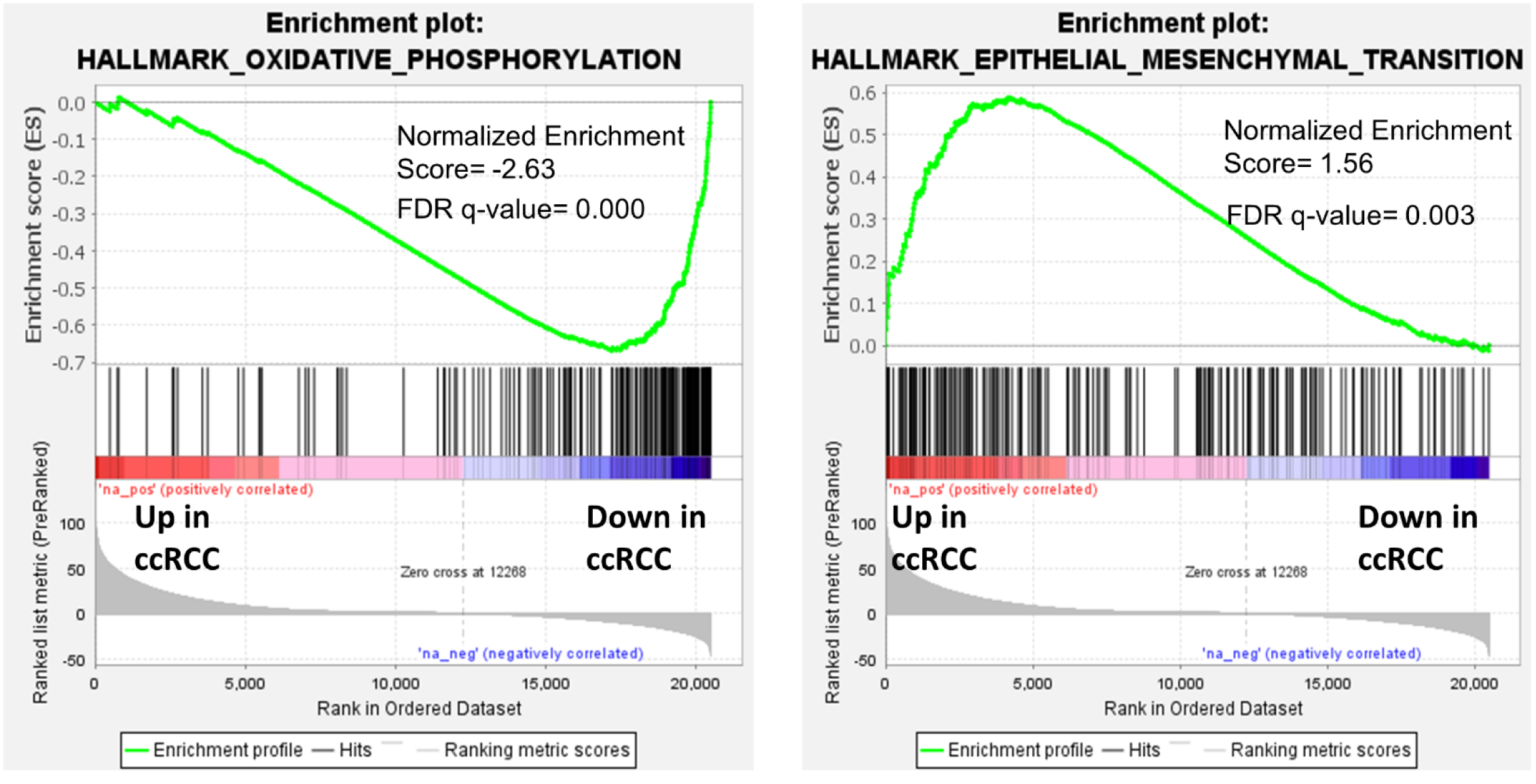
TCGA Pathway enrichment analysis reveals marked down-regulation of oxidative phosphorylation and upregulation of Epithelial mesenchymal transition in ccRCC.

**Fig S10.**
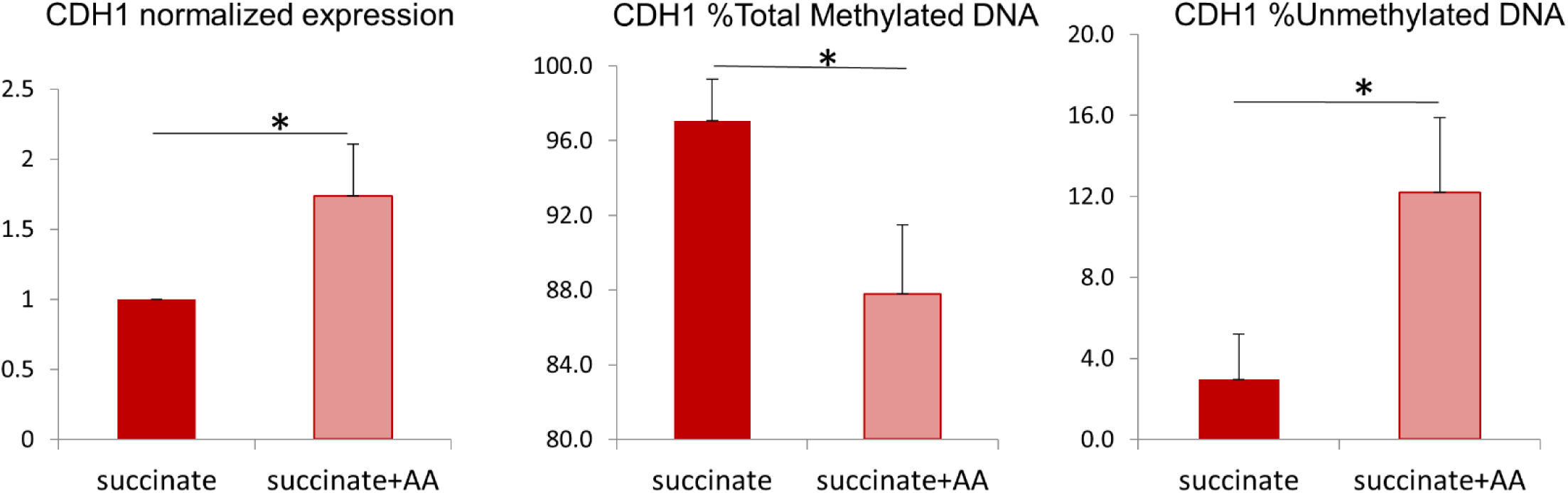
Addition of AA increases *CDH1* expression and decreases *CDH1* methylation in 7860 cells exposed to succinate.

**Fig S11.**
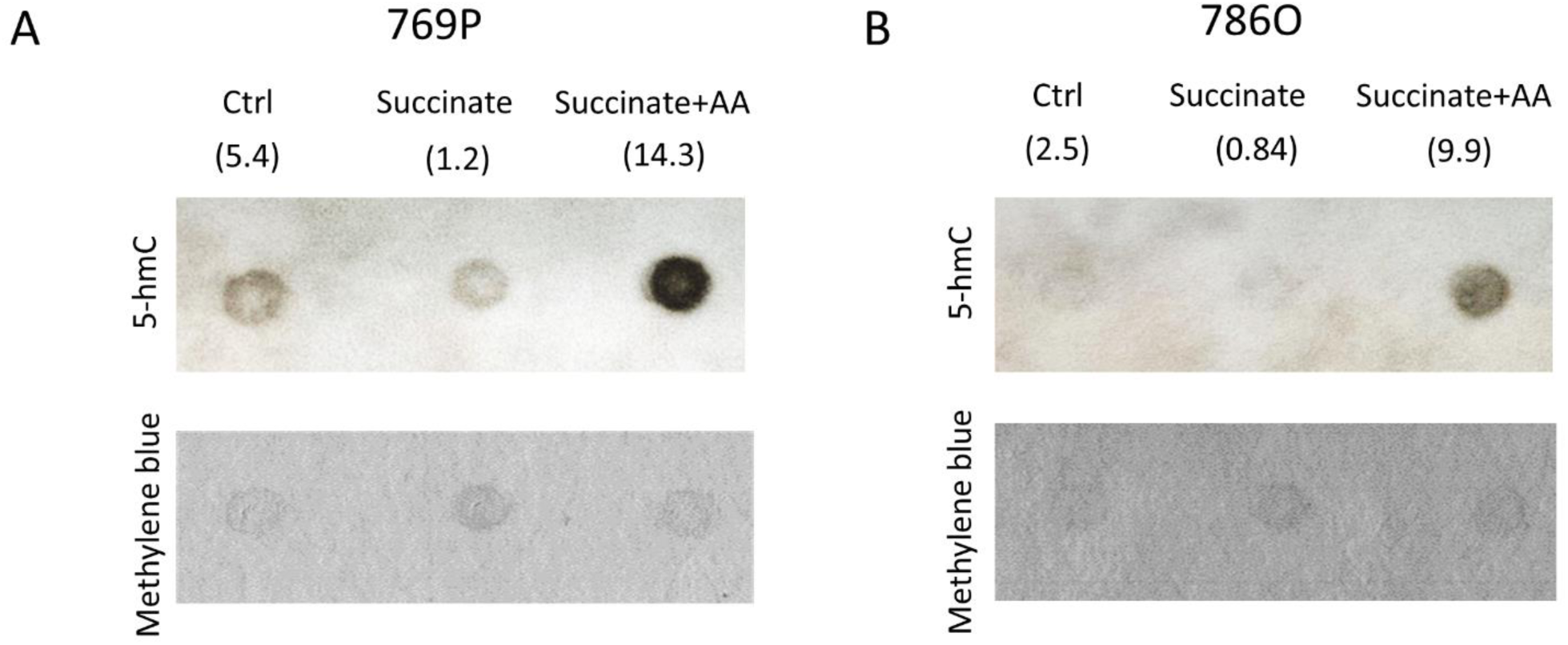
Succinate treatment of ccRCC cells decreases 5hmC levels whereas addition of AA markedly increases 5hmC levels (dot blot assay for 5hmC analysis with methylene blue as loading control)

**Fig S12.**
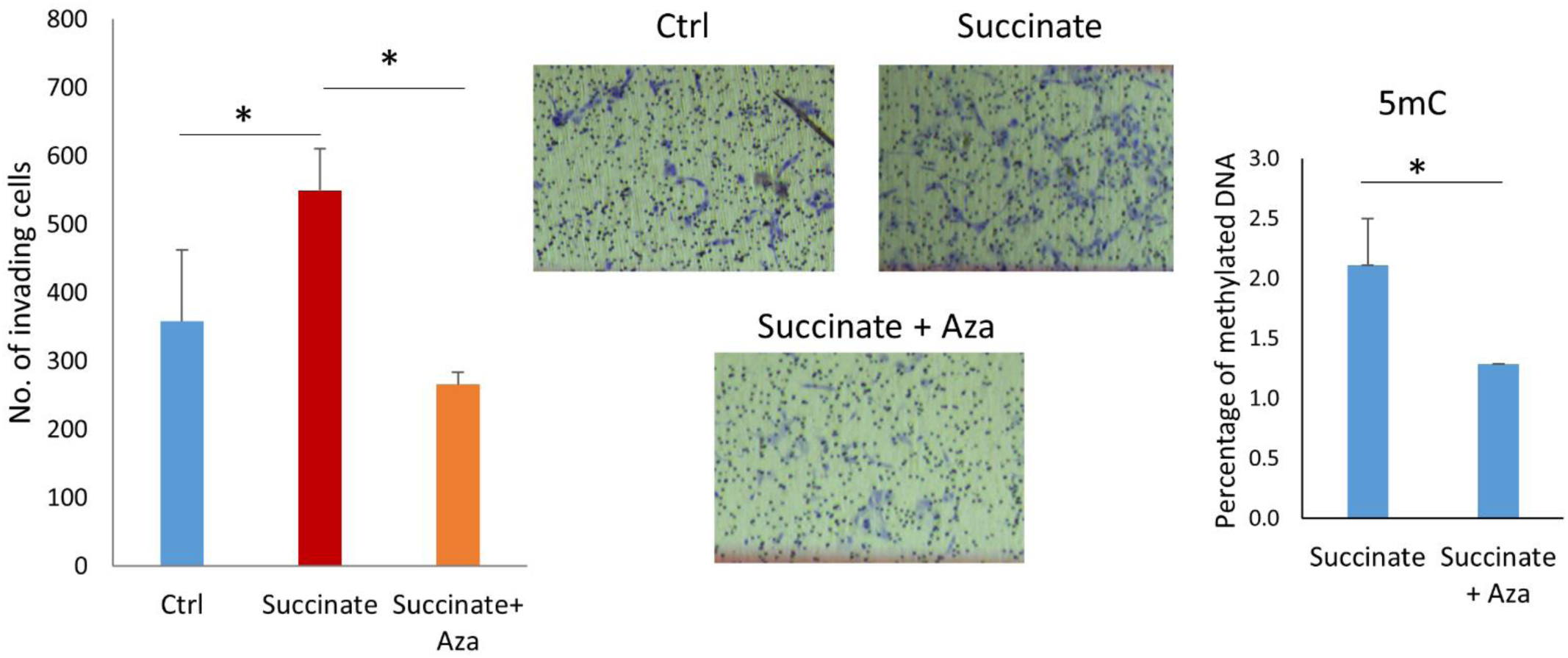
Succinate induced invasiveness of ccRCC cells is reversed by an archetypal DNA methyltransferase (DNMT1) inhibitor, Azacytidine.

**Table S1.** Top HIF2a binding sites annotated to miRNAs (+/-20kb from TSS) in ccRCC cell line 7860.

| Chr | Start | End | Score | Annotation | Distance To TSS | SYMBOL |
| --- | --- | --- | --- | --- | --- | --- |
| 11 | 565657 | 568457 | 200.8759 | Promoter (<=1kb) | 0 | MIR210HG |
| 22 | 26951178 | 26951289 | 151.3237 | Intron | 4232 | MIR548J |
| 9 | 1.27E+08 | 1.27E+08 | 85.53373 | Promoter (2-3kb) | 2873 | MIR181B2 |
| 17 | 75393066 | 75393136 | 64.21429 | Intron | 4771 | MIR4316 |
| 13 | 21007917 | 21007985 | 64.01366 | Intron | 13369 | MIR4499 |
| 4 | 7312177 | 7312251 | 57.36272 | Intron | 18566 | MIR4798 |
| 17 | 46709852 | 46709921 | 49.61469 | Distal Intergenic | -5385 | MIR196A1 |
| 1 | 2.08E+08 | 2.08E+08 | 46.43304 | Exon | -3346 | MIR29C |
| 3 | 1.34E+08 | 1.34E+08 | 41.49055 | Distal Intergenic | -10165 | MIR4788 |
| 22 | 20236657 | 20236734 | 41.38177 | Promoter (1-2kb) | 1700 | MIR1286 |
| 2 | 1.35E+08 | 1.35E+08 | 36.47723 | Distal Intergenic | -8460 | MIR3679 |
| 14 | 74225450 | 74225522 | 31.37908 | Promoter (1-2kb) | -1949 | MIR4505 |
| 17 | 25620936 | 25621022 | 30.56526 | Promoter (<=1kb) | 0 | MIR4522 |
| 22 | 35731633 | 35731751 | 30.19197 | Distal Intergenic | 15337 | MIR3909 |
| 12 | 1.05E+08 | 1.05E+08 | 27.30145 | Promoter (2-3kb) | -2797 | MIR3922 |
| 19 | 18392887 | 18392971 | 27.28812 | Promoter (<=1kb) | -56 | MIR3188 |
| 2 | 2.2E+08 | 2.2E+08 | 26.88108 | Exon | -4624 | MIR153-1 |
| 8 | 1.29E+08 | 1.29E+08 | 25.10832 | Intron | -14747 | MIR1205 |
| 21 | 26934457 | 26947480 | 22.74662 | Promoter (<=1kb) | 0 | MIR155HG |

Data accessed through GSE86092 <sup>5</sup>

## Conflict of interest statement

The authors declare no relevant conflict of interest.

## Abbreviations

ccRCC: clear cell renal cell carcinoma;
SDH: succinate dehydrogenase;
TET: Ten-Eleven Translocation;
AA: ascorbic acid

## Notes

### Competing Interest Statement

The authors have declared no competing interest.

### Summary of Updates

Title edit

